# Post-transfer instability, not initial transfer, limits dissemination of IncI1-*bla*_CTX-M-1_ plasmids between chicken and human *Escherichia coli*

**DOI:** 10.64898/2026.02.20.706986

**Authors:** M. Buffoni, E.A.J. Fischer, J. DelaFuente, A. San Millan, A.C. Schürch, R.J.L. Willems, J.A.G.M. de Visser

## Abstract

Plasmids are key vectors in the dissemination of antibiotic resistance genes. Within *Escherichia coli*, IncI1 plasmids carrying *bla*_CTX-M-1_ significantly contribute to extended-spectrum beta-lactamase (ESBL) resistance. To investigate their zoonotic dissemination potential, we compared the conjugative transfer rates, post-transfer stability, and short-term evolution of two IncI1-*bla*_CTX-M-1_ plasmids transferred from their original chicken *E. coli* host to a panel of chicken and human *E. coli*. *In vitro* conjugation assays revealed that transfer rates were not affected by recipient host origin or temperature and were primarily governed by recipient genotype. However, post-transfer plasmid loss rates were host-dependent, with four-fold higher loss rates in human than chicken-derived transconjugants. The total number of mutations accumulated post-acquisition was determined by the identity of the specific plasmid rather than the host origin, with each plasmid following a distinct evolutionary trajectory involving convergent parallel deletions. Specifically, one trajectory involved the loss of genes associated with plasmid stability, observed more frequently in human than in chicken strains, while the other involved the loss of part of the conjugation machinery. Our findings show that, although these plasmids transfer easily between bacteria from chicken to human hosts, they are less stable in their new host environment, which may restrict their overall spread. A comprehensive One Health risk assessment must consider these critical downstream dynamics of plasmid stability and host-specific evolution.

## Introduction

Antimicrobial resistance (AMR) poses a grave threat to modern medicine, with drug-resistant bacterial infections now ranking as a leading cause of death worldwide [1–3]. Within this crisis, the dissemination of Enterobacteriaceae resistant to third-generation cephalosporins (3GCs) is of particular concern [4]. The primary mechanism is the production of extended-spectrum beta-lactamases (ESBLs), most encoded by genes from the *bla*_CTX-M_ family [5, 6]. The global proliferation of these genes is largely attributable to their association with conjugative plasmids, which serve as vehicles for horizontal gene transfer (HGT) across diverse bacterial species and ecological niches [7, 8].

Antimicrobial resistance is a priority within the One Health framework; understanding the dynamics of plasmid-mediated transfer is essential to addressing how resistance moves between human, animal, and environmental sectors. [9]. Agricultural systems are a critical nexus for AMR transmission, where antibiotic use in livestock production enriches for resistant bacteria and the mobile genetic elements they carry [10, 11]. These resistance determinants are frequently shared between animal and human populations through various routes, including direct contact and the food chain [12].

Poultry production represents a significant interface for the exchange of resistance elements; molecular epidemiological studies have identified highly similar ESBL-carrying plasmids in *E. coli* isolated from poultry and human carriers, including occupationally exposed individuals and potential foodborne uropathogens [13,14]. The IncI1 plasmid group, often carrying the *bla*_CTX-M-1_ gene, is a paradigmatic example of this cross-host dissemination. Specific IncI1 plasmid sequence types, such as pMLST3 and pMLST7, are globally distributed in both agricultural and clinical settings, providing strong evidence for transmission [15, 16]. Although these narrow-host-range plasmids are specific for Enterobacteriaceae and depend on bacterial host factors for their persistence [17], their epidemiological success suggests they readily disseminate. Yet these epidemiological patterns do not illuminate the biological processes governing dissemination success. The presence of similar plasmids across different animal and human hosts can result from either the clonal expansion of resistant strains or the horizontal transfer of plasmids between diverse bacteria. Where HGT is the driver, this process begins with donor and recipient bacteria meeting and engaging in conjugation, followed by the challenge of plasmid persistence within the new population. Understanding the drivers of plasmid success in these scenarios requires the quantification of plasmid transfer rates between donor and recipient strains as a critical first step [18, 19], but should also involve determination of its subsequent fate, including its loss rate and fitness cost, as well as adaptation to the new host [20, 21].

To provide an integrated analysis of plasmid dissemination, stages often studied in isolation, we deconstructed the movement of two IncI1-*bla*_CTX-M-1_ plasmids, of pMLST3 and pMLST7, into human and chicken *E. coli*. We investigated: (i) the factors governing initial transfer; (ii) post-transfer stability; (iii) fitness effects; and (iv) the drivers of initial evolutionary changes, specifically distinguishing between adaptive responses (H1) and stress-induced mutagenesis (H2). We found that while conjugative transfer of these plasmids was largely unaffected by the host origin of recipient bacteria, their subsequent persistence within transconjugants was constrained by lower post-transfer stability and distinct, host-specific evolutionary adaptations.

## Results

### Plasmid transfer rates depend on recipient genotype, not host origin or temperature

To investigate transfer dynamics, we used two chicken-derived IncI1-*bla*_CTX-M-1_ plasmids (pMLST3 and pMLST7) representing major zoonotic lineages. These plasmids were selected as representative models for zoonotic spread because they possess a complete genetic toolkit for autonomous transfer, post-transfer stability, and multi-drug resistance (Fig. S1). Using the original wild-type chicken *E. coli* donors, we quantified conjugation rates across a diverse panel of 20 *E. coli* recipient strains (10 human, 10 chicken origin; Fig. 1A). The strains were dispersed across the *E. coli* phylogeny of >10,000 environmental and clinical isolates, ensuring a genomically diverse test panel [22]. Transfer rates were highly variable, spanning over six orders of magnitude across the different recipient strains. A nested ANOVA (Table S1) revealed no significant difference in transfer efficiency between the two plasmids (*F* = 0.485, *P* = 0.495). Consequently, data from both were analysed together for recipient effects. Furthermore, neither the recipient’s animal host origin (*F* = 0.319, *P* = 0.579) nor the animal host temperature (*F* = 0.010, *P* = 0.923) had a significant effect on the conjugation rate. The primary determinant of plasmid transfer was the specific genotype of the recipient strain (*F* = 11.90, *P* = 0.001), indicating that recipient-specific genomic makeup, rather than broad ecological origin or temperature, is the primary driver of plasmid spread.

**Figure 1:**
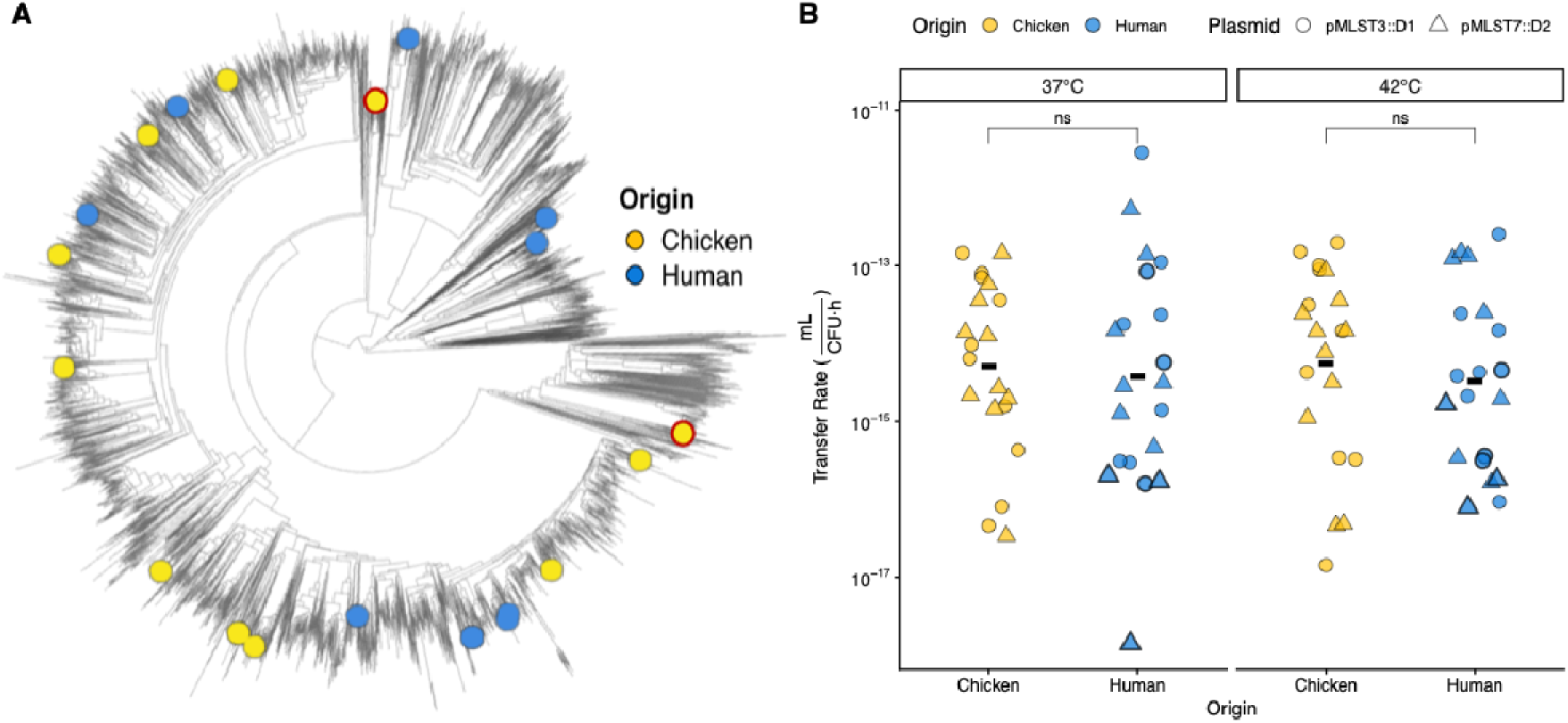
Genetic Diversity and Primary Transfer Dynamics. **(A)** Phylogenetic tree showing the genetic relationship of the two donors, indicated with red edges, and 20 recipient strains (10 human-origin, blue; 10 chicken-origin, yellow). The strains are placed within a reference collection of >10,000 *E. coli* genomes to illustrate the panel’s genetic diversity. (**B**) *In vitro* conjugation rates of the two chicken-derived plasmids (pMLST::*D*1, circle; pMLST::*D*2, triangle) into the recipient panel at 37°C and 42°C. Black horizontal bars indicate the mean transfer rate.

### Host-specific genomic makeup influences conjugation efficiency

Given the dominant role of the recipient strain and known dependence on donor-recipient relatedness [23], we investigated the influence of donor-recipient genomic relatedness and resident plasmid content on conjugation efficiency. Within the chicken host *E. coli* population, transfer rates were influenced by multiple genomic features. Accessory genome distance between the chicken donor and chicken recipient negatively affected conjugation rates (Spearman’s *ρ* = -0.47, adj. *P* = 0.005; Fig. 2A), as did the number of large resident plasmids (>10 kbp; *ρ* = -0.64, adj. *P* = 1.8 × 10□□; Fig. 2B). We quantified the abundance of functional defense and anti-defense systems on plasmids (Table S4). All systems, except two, were detected on large plasmids (>10 kb) in both human and chicken isolates. Chicken-associated recipients showed significantly greater defensive genomic investment in their plasmidomes than human strains (mean 3.2 vs. 1.2 systems per strain; Poisson GLM, *P* = 0.00376). In chickens, total system abundance was strongly negatively correlated with conjugation rates (*ρ* = −0.66, adj. *P* < 2.4 × 10□□; Fig. 2C). Consistent with this pattern, greater plasmidome divergence between donors and recipients was associated with higher transfer rates, especially in chicken (Fig. S2), whereas no correlation was observed with total or core genome distance (Fig. S3). These findings suggest that within a shared host environment, plasmids face ‘local rules of engagement’ governed by host-specific genomic barriers.

**Figure 2:**
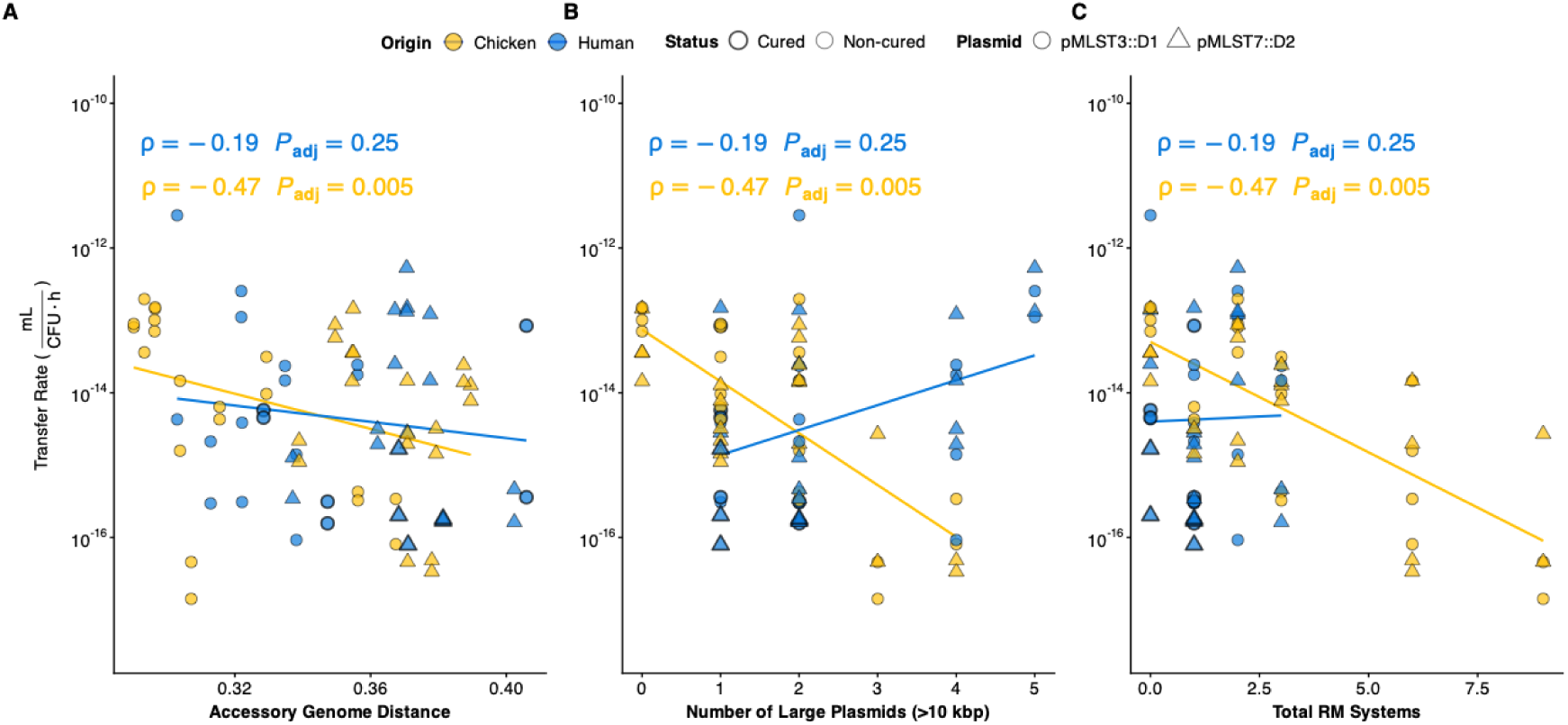
Host-Specific Genomic Barriers to Transfer. Association between plasmid transfer rates and the genomic makeup of recipient strains, stratified by host origin. Data points are shaped by donor plasmid type: pMLST3::D1 (circle) and pMLST7::D2 (triangle). (A) Relationship between transfer rate and the donor-recipietn accessory genome distance. (B) Relationship between transfer rate and the number of large resident plasmids (>10 kbp) in the recipient strain. (C) Relationship between transfer rate and the total count of functional defense and anti-defense systems/clusters identified in the recipient strain. Solid lines represent linear regressions per origin group. Spearman’s rank correlation (ρ) and Benjamini-Hochberg adjusted p-values are shown for each origin.

### Plasmid fitness costs are variable and trade-off against transmission at 42 °C

We next quantified the fitness effects of plasmid acquisition using maximum growth rate as a proxy, comparing transconjugants with their corresponding recipient strains. Plasmid carriage imposed a wide range of fitness effects, with relative growth rate values ranging from 0.49 to 1.19 across all conditions (Fig. S4). No significant difference in the overall average fitness cost was observed between human (mean = 0.926, SE = 0.02) and chicken (mean = 0.933, SE = 0.022) recipients (ANOVA, *F* = 0.888, *P* = 0.35). However, a significant interaction between donor plasmid identity and temperature was observed (ANOVA, *F* = 7.86, *P* = 0.007; Table S2): for plasmid pMLST3::D1, the fitness cost was higher at 42 °C, whereas it was lower for plasmid pMLST7::D2 (Wilcoxon signed-rank test, *V* = 164, *P* = 0.021; Fig. S4). We then tested for a transmission-cost relationship, specifically investigating whether higher transmission rates might impose a fitness cost on the host. Consistent with a transmission-virulence trade-off, faster transfer rates were associated with larger fitness costs for transconjugants in human strains at 42 °C (Spearman’s *ρ* = -0.67, adj. *P* = 0.006; Fig. S5). This suggests that high-stress temperatures, reaching the physiological limit for human hosts but typical for avian hosts, may create selective windows that favour plasmids with higher transmissibility despite the increased metabolic burden on the host.

### Plasmids exhibit high loss rates in transconjugants from a human origin

While transfer into recipients from human origin was not limited by host origin, we identified a significant post-transfer difference in plasmid maintenance. The average plasmid loss rate in human-derived transconjugants was approximately four-fold higher than in their chicken-derived counterparts (mean human = 0.97 h□¹, SE = 0.23; mean chicken = 0.22 h□¹, SE = 0.09; Wilcoxon rank-sum test, *V* = 328, *P* = 0.0002; Fig. 3A). This instability was not correlated with transfer rate, transconjugant relative growth rate, or genomic distance (Fig. S6). A nested ANOVA confirmed that recipient animal host origin was a significant factor explaining this variance (ANOVA, *F* = 34.46, *P* = 5.50 × 10□□), with variation among strains also contributing significantly to the observed instability (ANOVA, *F* = 3.54, *P* = 0.013; Table S3). While calculating loss-adjusted conjugation rates did not fundamentally alter the relative magnitudes of transfer rates (Fig. S7), it confirmed the primary conclusion of comparable net transfer between host origins (Fig. 3B). Established transconjugant fitness and fresh transconjugant loss rates were negatively associated for pMLST3::D1 (Spearman’s rho = -0.64, adj. *P* = 0.016; Fig. S8) but not pMLST7::D2 (rho = 0.11, adj. *P* = 0.680). Despite the mathematical interdependence of these variables and the finite strain panel, these distinct results suggest that pMLST3::D1 instability reflects a modulated, cost-dependent response to specific incompatibilities, while pMLST7::D2 instability is driven by a more generalized and disruptive stress response decoupled from immediate fitness effects.

**Figure 3:**
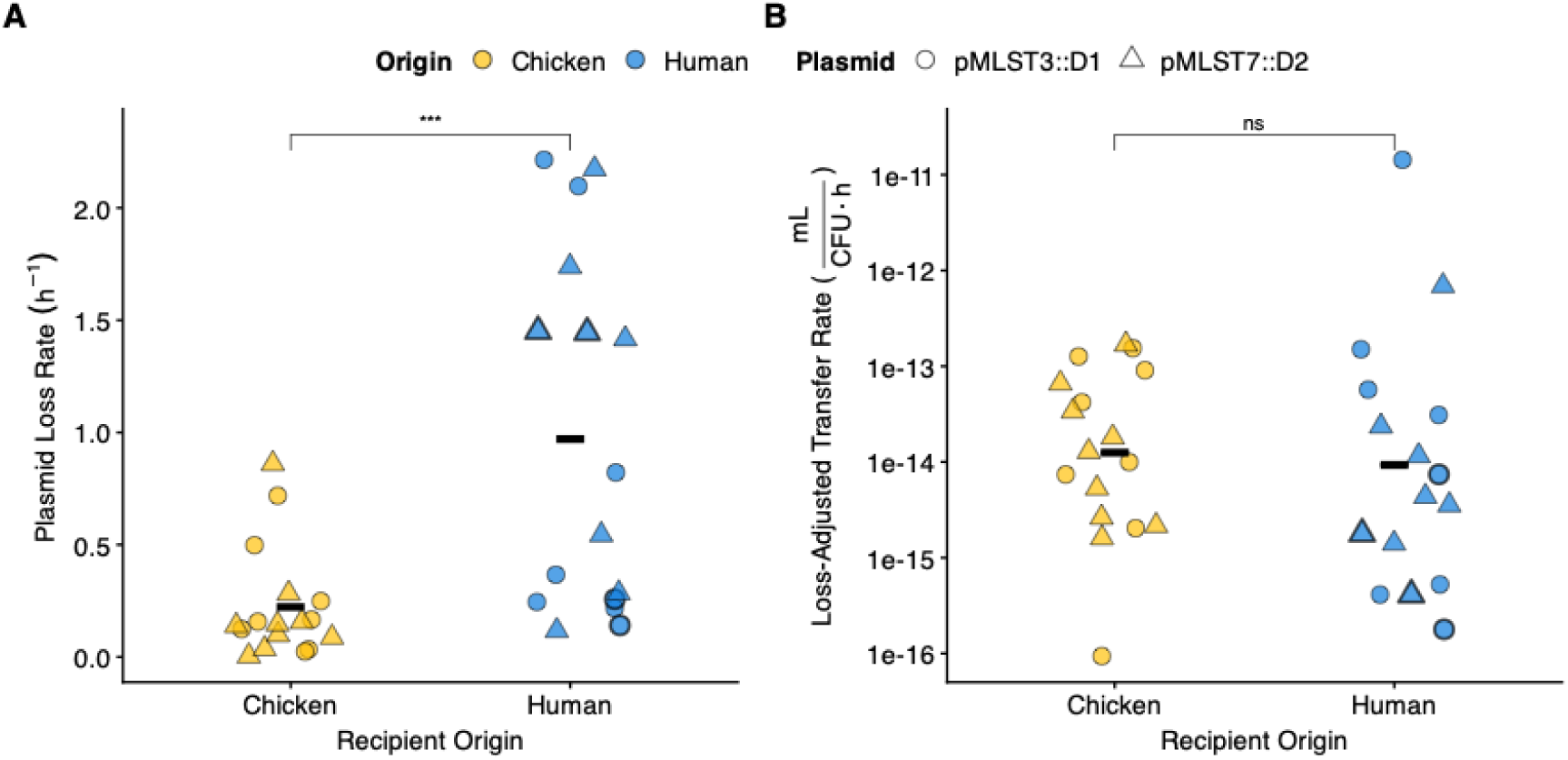
A Post-Transfer Barrier: Host-Origin Dictates Plasmid Stability. **(A)** Plasmid loss rates. Rates were calculated as the fraction of plasmid-carrying cells lost per hour of growth in antibiotic-free media (37°C), determined via differential plating. Human-derived transconjugants (Ts) exhibited significantly higher plasmid instability compared to chicken-derived strains (Wilcoxon test, *P* < 0.001). **(B)** Loss-adjusted transfer rates. To determine the effective dissemination potential, initial transfer rates were adjusted for post-transfer loss. While initial transfer was efficient, the resulting stability-adjusted rates in human hosts illustrate the impact of host-dependent persistence (Wilcoxon test, *P* = ns). Black horizontal bars indicate the mean. Bold borders highlight plasmid-cured strains.

### Post-conjugational evolution is shaped by host environment and plasmid identity

We were then interested to see whether plasmid acquisition triggered immediate genetic change in the genome of the transconjugants that could facilitate rapid adaptation. To identify mutations specifically linked to the period following plasmid acquisition, we compared the whole-genome sequences of the transconjugants directly to their respective parental recipient strains. Mutational analysis was performed on 31 transconjugant lineages following the conjugation event and approximately 60 generations of subsequent growth. Mutation frequencies were highly variable: 5 had no mutations, while the remaining 26 accumulated at least one, with the highest number observed being 39 mutations and with an average number of mutations of 5.56 in chicken strains and 6.57 in human strains (Fig. 4A, Wilcoxon *p* > 0.05).

**Figure 4.**
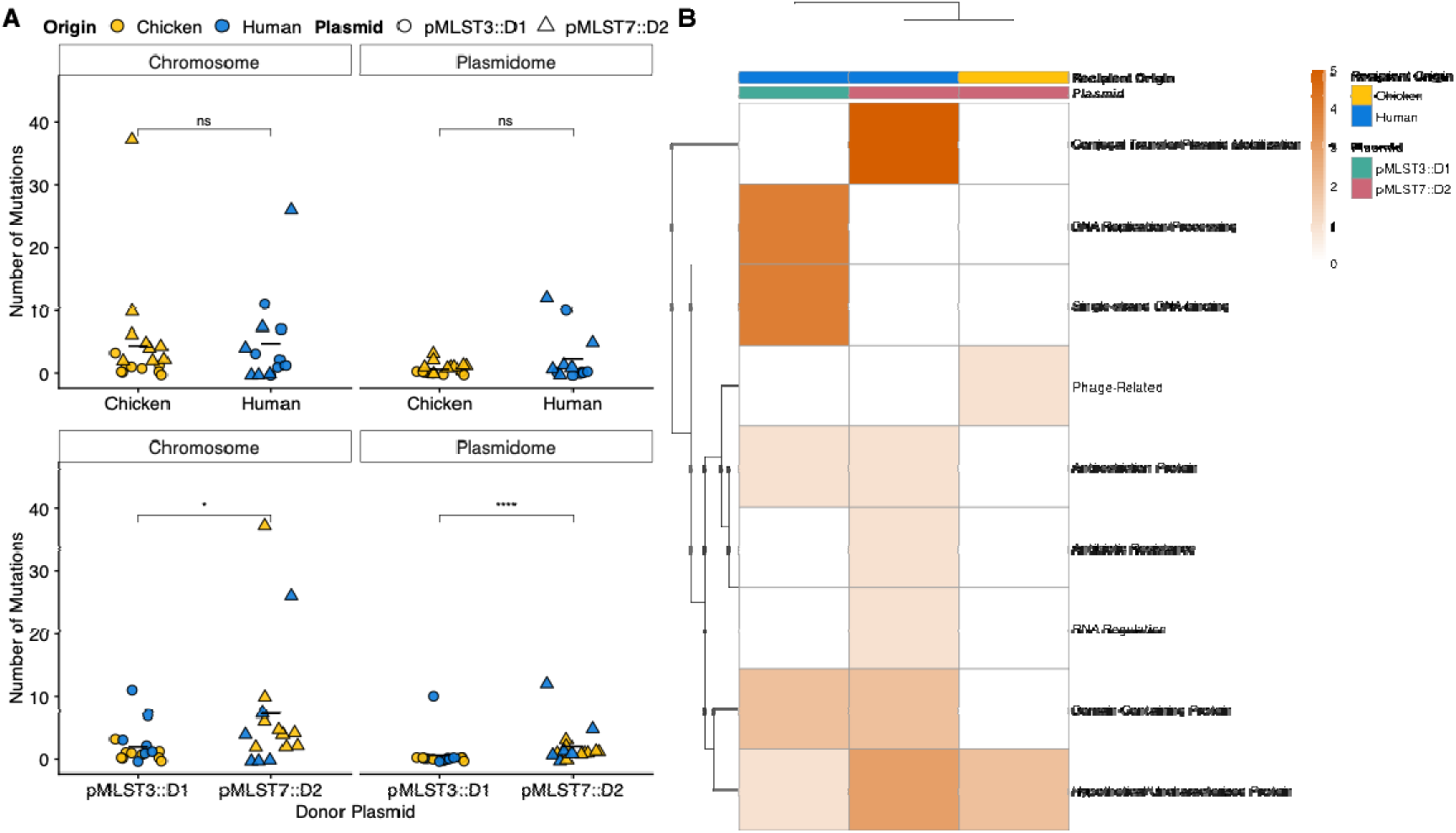
Host-Specific Mutational Response Following Plasmid Acquisition. **(A) Quantification of mutations.** Scatter plots display the total mutation count across the chromosome and plasmidome, stratified by recipient origin (top) and specific plasmid identity (bottom). Points are colored by recipient origin (yellow: chicken; blue: human) and shaped by plasmid type (circles: pMLST3::D1; triangles: pMLST7::D2). Horizontal black bars indicate the mean. **(B) Functional landscape of plasmidome mutations.** Clustered heatmap showing mutation counts across functional gene categories. Clustering analysis reveals distinct evolutionary trajectories dictated by the interaction between plasmid identity and host environment. The intensity of orange shading corresponds to the total mutation count within each category. Statistical significance (Wilcoxon test): ns, *P* > 0.05; *, *P* < 0.05; ****, *P* < 0.0001).

The correlation between mutation frequencies, relative fitness, and donor-recipient genomic distance varied markedly by donor plasmid and recipient host origin. In pMLST3::*D*1-harboring transconjugants, mutation frequencies correlated positively with genomic distance (Spearman’s *ρ* = 0.58, adj. *P* = 0.035) and negatively with relative fitness (ρ = –0.67, *adj. P* = 0.009). Similarly, chicken-origin transconjugants showed strong positive correlations with genomic distance (ρ = 0.86, adj. *P* < 0.001) and negative correlations with fitness (ρ = –0.58, adj. *P* = 0.022). In contrast, no significant correlations were observed for pMLST7::D2-harboring or human-origin transconjugants (Fig. S8A–D) with fitness and genomic distance. However, transconjugants that acquired the pMLST7::D2 plasmid showed significantly more mutations in both the chromosome and plasmidome compared to those carrying the pMLST3::D1 plasmid (Fig. 4B). Lastly, mutation frequencies were unrelated to plasmidome similarity or loss rate across all analyses (data not shown). These results indicate that both plasmid type and animal host origin shape distinct patterns of post-conjugation mutation accumulation and associated fitness effects.

Quantification of these mutations showed that while the central tendency (median) of total chromosomal and plasmidome mutations was variable across lineages (Wilcoxon test, ns; Fig. 4A), the host origin significantly impacted the mutational rate and target landscape. In the plasmidome, human-derived transconjugants exhibited a significantly higher mutational rate compared to chicken-derived transconjugants (Poisson GLM, *P* = 1.6×10^−4^). This was driven by high-intensity mutational activity in specific human backgrounds, averaging 2.31 mutations per lineage, whereas chicken lineages remained largely stable (0.61 mutations per lineage). Furthermore, while the total chromosomal mutation counts were comparable between hosts, the frequency of mutations within specific functional categories was host biased. Human-origin transconjugants accumulated a significantly higher rate of mutations in regulatory proteins (Poisson GLM, *P* = 0.022) and phage/transposable elements (Poisson GLM, *P* = 0.025) compared to chicken-origin transconjugants (Fig. S10). This suggests an animal host-specific response to conjugation, particularly impacting mobile genetic elements and regulatory networks in the chromosome of human strains, and affecting the plasmidome in human-derived transconjugants.

Most notably, we found a host-specific response for the two donor plasmids, primarily through large parallel deletions (Fig. 5A). The parallel nature of these deletions was suggestive of an adaptive evolutionary response. In pMLST3::D1, deletions occurred within a region containing stress-response genes, including the SOS-inhibition genes *psiA* and *psiB*, in 6 of 9 human and 1 of 9 chicken transconjugants (Table S5). In pMLST7::D2, parallel deletions predominantly involved the conjugation machinery (*trbA/B*, *nikB*) and specific toxin-antitoxin (TA) components (*sok*, *pndC*) across 8 of the 16 analysed transconjugants (Table S6). The absence of insertion sequences flanking these common deletion breakpoints is consistent with selection rather than structural predisposition, strongly indicating an adaptive evolutionary response for these parallel deletions. Beyond these parallel events, we observed a diverse array of lineage-specific deletions unique to individual transconjugants (Table S7). In pMLST3::D1, the extreme instability observed in specific human lineages was associated with extensive deletions that encompassed essential maintenance machinery, including the *parM/parB* partitioning and *relE/impCAB* TA systems (Table S8). For pMLST3::D1, a greater reduction in plasmid size significantly correlated with both higher fitness costs and greater plasmid instability (Fig. 5B). Whether incompatibility drove these deletions or genomic loss exacerbated instability remains undetermined; however, the most incompatible pairs clearly underwent the most extensive remodelling. The significant correlation in pMLST3::D1 is likely explained by the fact that only the largest lineage-specific deletions resulted in the loss of partitioning and TA systems. In contrast, no such correlation was observed for pMLST7::D2, where TA components (*sok, pndC*) were already lost in parallel across many lineages while the partitioning system was largely preserved, potentially decoupling total deletion size from plasmid persistence.

**Figure 5:**
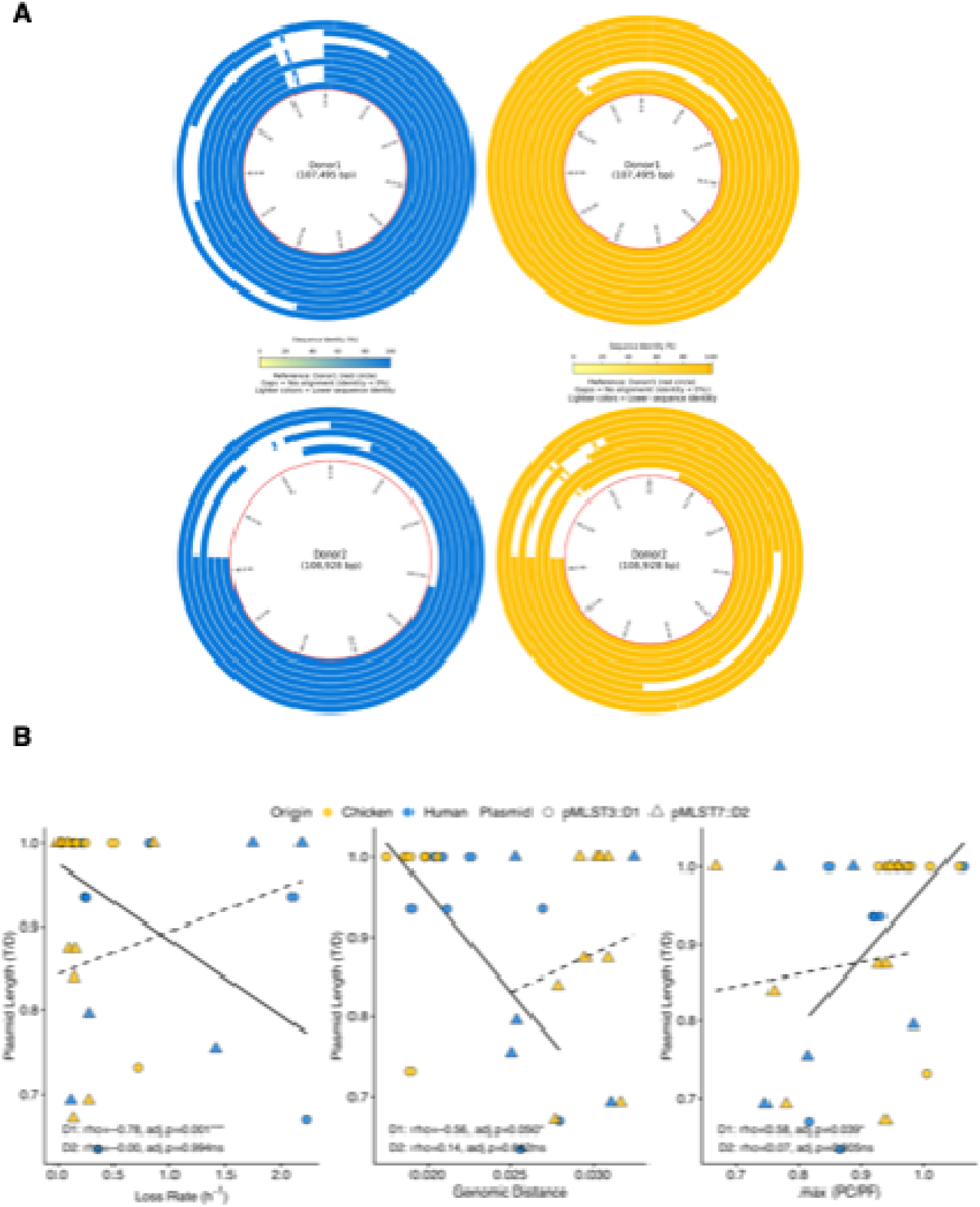
Divergent parallel Deletions in New Hosts. **(A)** Circular genome alignment plots showing deletions in donor plasmids after transfer into human (blue) and chicken (yellow) backgrounds. The inner most red circle is the *D* reference sequence. Gaps in the concentric circles (representing independent *T*s) indicate deletions. (**B**) Correlation between the relative length of the evolved plasmid pMLST3::*D*1 (solid line) and pMLST7::*D*2 (dashed line) and three key parameters. Points are colored by host origin and shaped by *D* plasmid.

### Prior host exposure mitigates fitness costs but not plasmid instability

Finally, we investigated whether a history of plasmid-harboured *bla*_CTX-M-1_ gene carriage affects the dynamics of subsequent plasmid acquisition in human recipients. Three of the ten human recipients were strains that originally and naturally carried plasmids (two IncI1 and one IncFII) harbouring the *bla*_CTX-M-1_ gene, the same resistance marker present on our donor plasmids (pMLST3::D1 and pMLST7::D2). Before the conjugation assays, we performed a curing step to remove these resident *bla*_CTX-M-1_ plasmids. Genomic analysis confirmed the successful removal of the target plasmids, which ranged from moderately related IncI1 types to a distantly related IncFII type (Table S8). Notably, this analysis also revealed that the curing process induced collateral changes in the non-target resident plasmidome, including plasmid loss and rearrangements (Table S9). However, these alterations involved mostly small plasmids and may partly reflect artifacts inherent to long-read sequencing limitations.

This human subgroup selection allowed us to evaluate whether a host already ‘familiar’ with the metabolic and regulatory demands of this specific resistance cargo and similar plasmid backbones would better tolerate new acquisitions. We compared these three cured strains to the seven ‘naive’ human strains (strains never exposed to *bla*_CTX-M-1_).

This history of carriage had plasmid-specific effects on transfer: it significantly reduced the conjugation rate for pMLST7::D2 into cured hosts compared to naive hosts but had no significant effect on pMLST3::D1 (Fig. 6A). Crucially, we observed that previous exposure significantly reduced the metabolic burden of new plasmid acquisition. We calculated the relative growth rates of transconjugants derived from cured versus naive hosts. For both donor plasmids, the cost of acquisition was significantly lower in cured strains than in naive strains (Fig. 6B). Cured strains carrying a new donor plasmid showed average relative growth rates of 0.983 (pMLST3::D1) and 1.089 (pMLST7::D2), which were significantly higher than the 0.922 (pMLST3::D1) and 0.844 (pMLST7::D2) observed in naive transconjugants.

**Figure 6:**
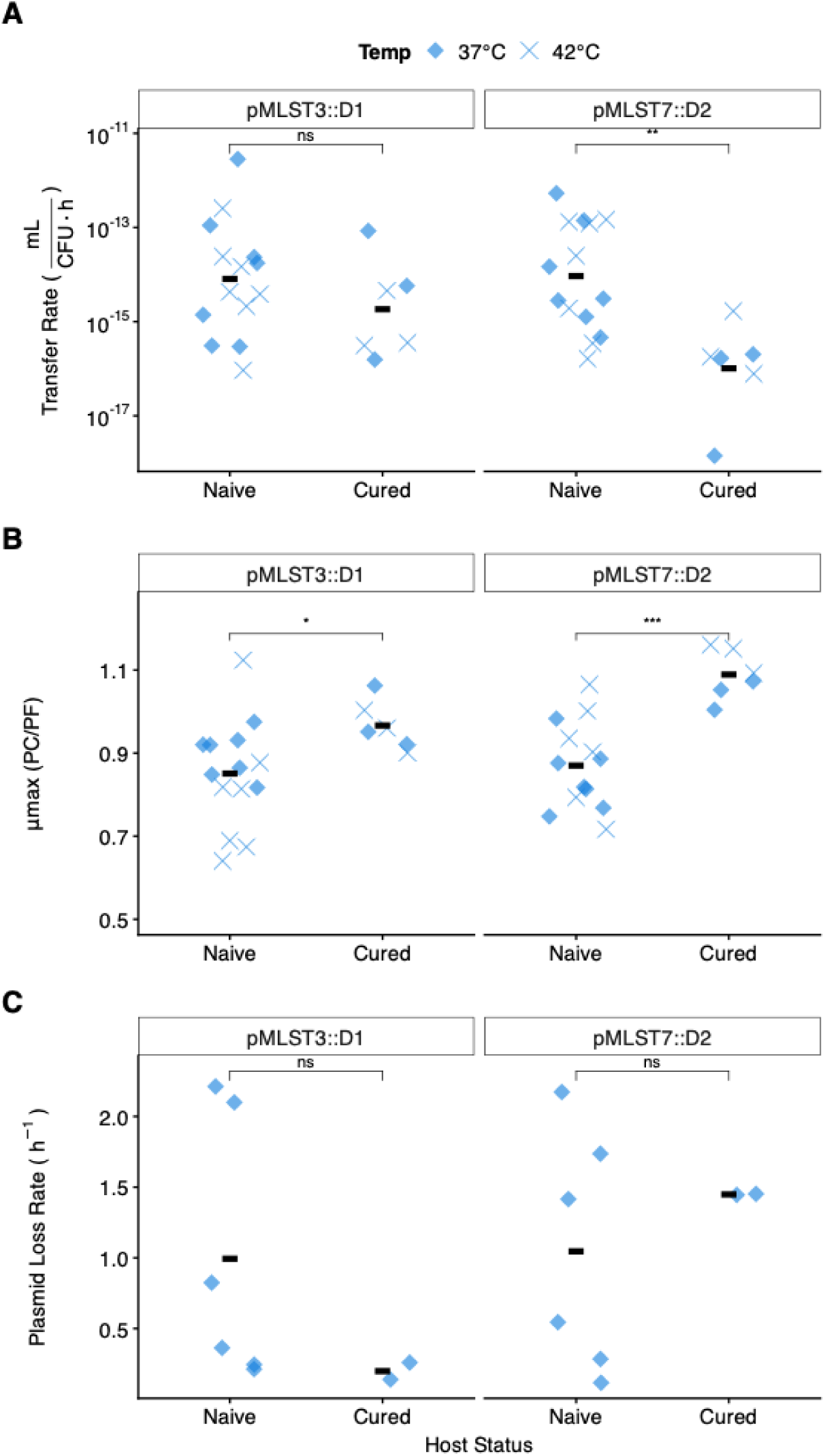
Effect of prior host exposure to plasmids carrying *bla*_CTX-M-1_ on plasmid transfer dynamics. A comparison of key traits for plasmids pMLST3::*D*1 and pMLST7::*D*2 following transfer into either a naive *R* host or a host previously cured of a native *bla*_CTX-M-1_ plasmid. The three panels display (**A**) the conjugation transfer rate, (**B**) the relative growth rate of the resulting *T*s (plasmid-carrying / plasmid-free), and (**C**) the plasmid loss rate. Measurements were conducted at both 37°C (blue diamonds) and 42°C (light blue crosses). Within each plot, individual points represent biological replicates, and black horizontal lines denote the group median. Statistical comparisons between naive and cured host statuses were performed using the Wilcoxon rank-sum test, with significance indicated as ns (not significant), * (p < 0.05), ** (p < 0.01), or *** (p < 0.001).

However, this fitness benefit did not result in improved plasmid maintenance; loss rates in the cured strains remained high and were not significantly different from those in naive human strains (Fig. 6C). This suggests that while previous carriage of a *bla*_CTX-M-1_ plasmid may mitigate the general metabolic burden of carriage, it does not resolve specific vertical incompatibility issues between the donor plasmids and the human host background.

## Discussion

The dissemination of antibiotic resistance plasmids from animal reservoirs to human pathogens is a multi-stage process. We deconstructed this process for the dissemination of two common conjugative IncI1plasmids among *Escherichia coli* isolates from chicken and humans, by analysing the distinct phases of conjugative transfer, post-transfer persistence, and subsequent rapid evolutionary adaptation occurring within approximately 60 generations post-acquisition. Our main finding indicate that the initial entry of these plasmids is not restricted by host origin or temperature while their long-term persistence is challenged by host-specific incompatibilities.

We found that neither host origin nor physiological temperature were primary determinants of plasmid transfer. Instead, transfer rates were strongly affected by recipient genotype, with a significant effect of donor - recipient accessory genome distance, especially the presence of large plasmids, in chicken-derived recipients. Our findings highlight the role of resident big plasmids as primary reservoirs for defence and anti-defence systems, functioning as gatekeepers of HGT. The ‘defensive enrichment’ observed in chicken recipients suggests that these resident plasmids provide a robust barrier against the invasion of new genetic material. Conversely, the lower prevalence of these systems in human isolates may reflect a more permissive landscape for plasmid acquisition. This suggests that each community has evolved host-specific mechanisms, such as restriction-modification (RM) and CRISPR-Cas systems [24, 25], to regulate horizontal gene flow.

This defensive disparity likely reflects the heightened phage pressure of chicken bedding compared to the anaerobic environment of the human gut. Beyond viral defence, these systems frequently mediate inter-plasmid conflict. Plasmid-borne CRISPR-Cas loci, particularly Type IV-A3 systems, are widespread and primarily target other mobile genetic elements rather than viruses, suggesting their role in “inter-plasmid warfare” [26, 27]. Furthermore, by encoding methyltransferases and anti-restriction genes, plasmids can counteract host RM systems, thereby actively shape conjugation dynamics and reduce the host’s effectiveness as a barrier to gene flow [28].

While these specialized defences govern the initial rules of engagement during entry, the subsequent challenge shifts to maintenance. Following successful transfer, we identified a four-fold higher rate of plasmid loss in human relative to chicken transconjugants, indicating a profound stability challenge. Such instability is consistent with known outcomes for plasmids in novel bacterial hosts, where mismatched replication control, partitioning systems and toxin-antitoxin dependencies can lead to rapid loss during cell division [29,30]. Furthermore, the fitness consequences of plasmid carriage varied significantly with plasmid identity, temperature, and recipient host origin. The opposing effects of temperature on plasmid fitness costs and the transmission-cost trade-off observed only in human strains at 42°C suggest a complex interplay. While 42 °C is physiological for avian hosts, it represents a severe stressor for human-derived *E. coli*. The entry of conjugative plasmids might trigger a host SOS response [31], which, in combination with thermal stress, could potentially exacerbate the metabolic burden on the host [32]. This cumulative stress might explain why the transmission-cost trade-off was observed specifically in human backgrounds at 42 °C.

The observed host-induced stress and subsequent evolutionary dynamics are likely mechanistically linked: stress-induced mutagenesis can increase the local mutation rate, generating the raw mutational substrate (e.g., via error-prone polymerases or replication errors) upon which natural selection may then act, favouring mutations (including deletions) that confer an immediate fitness. The generalized stress response in human-derived recipients likely reflects plasmid–host incompatibility rather than a uniquely human trait, and similar mutagenic effects may arise in any mismatched host–plasmid combination. Nevertheless, the human host environment appeared to trigger a particularly pronounced stress response, as indicated by a mutational pattern uncorrelated with genetic distance and an excess of mutations within the transconjugants own resident plasmids. While stress-induced mutations are non-adaptive byproducts of this process, selection may act upon them. Notably, they were enriched in genes encoding phage-related proteins and transposons (known hotspots for genomic instability) and in global regulatory proteins, the latter being consistent with systemic adaptation to stress [33]. In chickens, by contrast, the mutational response appears more tightly coupled to the genetic distance between donor and recipient strains. A similar pattern is seen between the donor plasmids themselves: for pMLST3::*D*1, the number of mutations correlates with genetic distance, whereas for pMLST7::*D*2, this relationship is lost. This likely reflects the differing ranges of genetic divergence among their respective donor *E. coli* backgrounds, pMLST3::*D*1 being transferred across recipients spanning a broad spectrum from closely related to more distant genotypes, while pMLST7::*D*2 was transferred primarily among more distantly related recipients. These contrasting mutational landscapes, evident both across host origins and between the donor plasmids, suggest distinct underlying dynamics. The generalized stress response observed in human *E. coli* and with pMLST7::*D*2 likely arises when plasmid–host interactions constitute a disruptive or novel challenge, overwhelming established adaptive systems. Conversely, the modulated, incompatibility-dependent response seen in chicken *E. coli* and with pMLST3::*D*1 implies a degree of prior co-adaptation or limited genetic divergence, allowing hosts or plasmids to mount a more proportional, finely tuned response to incompatibility.

This stressful environment posed by the new plasmid-host combinations in transconjugants triggered different adaptations of the two plasmids. Plasmid pMLST7::*D*2 shed its universally costly conjugation machinery in 50% of the transconjugants, a known compensatory mechanism [34, 35]. In contrast, the deletions in pMLST3::*D*1 highlight a fundamental evolutionary trade-off: short-term survival under selection versus long-term plasmid stability.

The idea that short-term adaptation can incur long-term costs is a well-documented evolutionary phenomenon [36, 37]. The positive correlation we observed between relative deletion size and fitness cost suggests that these genomic losses may reflect: (i) costly deletions caused by stress-induced mutagenesis in “problematic” host-plasmid combinations; (ii) incomplete adaptations, where larger deletions are required to survive in combinations imposing higher initial costs; or (iii) the targeted shedding of anti-defence systems that become redundant in a novel host landscape.

The loss of the ‘leading region’, a specialized genomic compartment recently shown to encode a high diversity of anti-defence systems [38], specifically the SOS-inhibitors *psiA* and *psiB*, is a key example of this third possibility. While these systems are likely essential for navigating the ‘defensive enrichment’ we identified in chicken isolates, they might represent redundant metabolic baggage in the defence-poor environment of human *E. coli*. The lack of selective pressure to maintain these anti-restriction modules in human hosts likely drives their rapid deletion to alleviate fitness costs.

However, this adaptation represents a *de facto* Pyrrhic victory. In the process of shedding redundant genes to reduce metabolic burden, the plasmid inadvertently discards co-located partitioning systems or toxin-antitoxin modules, often termed ‘addiction systems’, that are crucial for its own segregational stability [39]. What initially appears to be an adaptive refinement in the human host compromises long-term persistence, rendering the plasmid less capable of remaining within the population once selective pressure is removed.

Our analyses of three human recipient strains cured of their native *bla*_CTX-M-1_ plasmids revealed a decoupling between plasmid fitness cost and stability. Prior exposure to these resistance plasmids reduced the metabolic burden of a newly acquired plasmid but did not prevent segregation loss during cell division. This indicates that while host compensatory mutations can mitigate fitness costs [40, 41], they do not necessarily resolve incompatibilities with an incoming plasmid’s replication or partitioning systems, thereby limiting plasmid dissemination across host populations [42]. This fitness improvement likely reflects the re-establishment of a plasmid–chromosome crosstalk that had been previously optimized in the host that could persist after plasmid curing [43]. The lower plasmid-associated cost in strains naturally carrying the plasmid than in naïve strains suggests that these adaptations pre-condition hosts to better tolerate the metabolic demands of the tested large plasmids [44]. However, the apparent reduction in transfer rate may reflect efficient plasmid entry followed by rapid post-transfer loss, illustrating how endpoint conjugation assays can obscure differences between plasmid entry and stable maintenance [45].

In conclusion, our findings demonstrate that while chicken-derived IncI1-*bla*_CTX-M-1_ plasmids can efficiently transfer to human *E. coli*, their subsequent persistence is severely limited by high instability. A comprehensive One Health risk assessment must therefore move beyond simple transfer rate measurements and account for these critical downstream events of plasmid stability and host-specific evolution that ultimately govern the success of ESBL plasmid dissemination from animal reservoirs.

## Methods

### Bacterial Strains, Plasmids, and Culture Conditions

The study was designed around two chicken-derived *Escherichia coli* donor strains and a recipient panel comprising 20 antibiotic-susceptible *E. coli* isolates (10 from healthy human faeces, 10 from healthy commercial chicken faeces) (Table 1). *D*1 (strain BR4) harboured an IncI1-pMLST ST3 plasmid carrying the *bla*_CTX-M-1_ gene (107,495 bp). *D*2 (strain 38.52) harboured an IncI1-pMLST ST7 plasmid, also carrying *bla*_CTX-M-1_ (108,928 bp) (Table S7). A whole-plasmid Mash comparison (v2.2.2) yielded a genetic distance of 0.00763, confirming the plasmids are highly related. To enable selection of transconjugants, all recipient strains were rendered resistant to chloramphenicol (CHL; 32 µg/mL) by introducing the non-mobilizable plasmid pBGC [44]. Finally, three human recipient strains (WT4, WT5, WT8) were cured of their native *bla*_CTX-M-1_ plasmids using the CRISPR-Cas9-based system described below [47] (graphical maps of plasmids used for curing and selection are provided in Figure S11). All bacterial strains were routinely cultured in Lysogeny Broth (LB) at 37°C.

**Table 1.** List of *Escherichia coli* strains used in the study.

| Strain Role | Strain ID | Host Origin | Isolation Year | Plasmid-Cured | Source/ Reference |
| --- | --- | --- | --- | --- | --- |
| Donor (D1) | BR4 | Chicken | 2023 | No | This study |
| Donor (D2) | 38.52 | Chicken | 2014 | No | De Been et al. [8] |
| Recipient | MH1 | Human | 2023 | No | This study |
| Recipient | JH2 | Human | 2023 | No | This study |
| Recipient | 3H1 | Human | 2023 | No | This study |
| Recipient | 3H4 | Human | 2023 | No | This study |
| Recipient | 4H1 | Human | 2023 | No | This study |
| Recipient | 5H1 | Human | 2023 | No | This study |
| Recipient | 5H4 | Human | 2023 | No | This study |
| Recipient | WT4 | Human | 2017 | Yes ☐ | Dorado-Garcia et al. [57] |
| Recipient | WT5 | Human | 2017 | Yes ☐ | Dorado-Garcia et al. [57] |
| Recipient | WT8 | Human | 2017 | Yes ☐ | Dorado-Garcia et al. [57] |
| Recipient | 1B1 | Chicken | 2023 | No | This study |
| Recipient | 1B4 | Chicken | 2023 | No | This study |
| Recipient | 1B6 | Chicken | 2023 | No | This study |
| Recipient | 1B8 | Chicken | 2023 | No | This study |
| Recipient | 2B1 | Chicken | 2023 | No | This study |
| Recipient | 2B7 | Chicken | 2023 | No | This study |
| Recipient | 3B6 | Chicken | 2023 | No | This study |
| Recipient | 4B1 | Chicken | 2023 | No | This study |
| Recipient | 4B3 | Chicken | 2023 | No | This study |
| Recipient | 5B1 | Chicken | 2023 | No | This study |

☐ Strains were cured of their native *bla*<sub>CTX-M-1</sub>-harboring plasmid.

### Plasmid Curing

A single-guide RNA (sgRNA) targeting a conserved region of the *bla*_CTX-M_ gene was introduced into the pLC10 vector via Golden Gate assembly (New England Biolabs) [48]. Human isolates WT4, WT5, and WT8 were cured using the apramycin-resistant variant of this CRISPR-Cas9 system, pLC10-Apra [47]. Plasmids were introduced via electroporation using 0.1 cm cuvettes and a 1.8 kV pulse (MicroPulser Electroporator; Bio-Rad Laboratories). Transformants were selected on apramycin (30 mg/L) and verified by PCR. Curing was induced by resuspending colonies in LB broth supplemented with 0.2 mg/L anhydrotetracycline and 0.08 mM isopropyl β-D-1-thiogalactopyranoside (IPTG). After a 2-hour incubation at 30 °C with agitation (250 rpm), cells were streaked onto LB agar and incubated overnight at 37 °C; this temperature shift facilitated the loss of the curing vector via its thermosensitive pSC101 origin of replication. Colonies sensitive to both apramycin and cefotaxime (CTX), the original plasmid-associated antibiotic, were recovered and confirmed as cured through PCR and Oxford Nanopore whole-genome sequencing.

### Monoculture growth assay

To assess plasmid fitness effects and provide parameters for conjugation models, monoculture growth assays were performed for donor, recipient, and transconjugant strains. Cultures were initiated from glycerol stocks in 3 mL LB; tranconjugants strains were supplemented with CTX (1 µg/mL) to ensure 100% plasmid carriage at the onset. After overnight incubation at 37 °C and agitation (250 rpm), cultures were diluted 1:100 in antibiotic-free LB and grown to mid-exponential phase (OD□□□ ≈ 0.15). Each was then diluted to ∼10□ CFU mL□¹ in 150 µL LB and dispensed in triplicate into 96-well plates with a 50 µL mineral oil overlay. Plates were incubated in a Victor³ plate reader (PerkinElmer) at 37 °C or 42 °C under static conditions, with OD□□□ readings every 6 min for 24 h. Maximum growth rates (μ*_max_*) were estimated using the *all_linear* method in the R package *growthrates* (v0.8.4) [49]. Relative fitness was expressed as the ratio of the μ*_max_* of the transconjugant to its corresponding isogenic recipient.

### Conjugation Rate Measurement

Donor and recipient strains were grown from -80°C stocks to mid-exponential phase (OD600 ≈ 0.15) in LB. Exponential phase cultures of donor and recipient strains were mixed 1:1 (v/v) to initiate mating. Immediately upon mixing (*t* = 0) and after four hours (*t* = 4) at either 37°C or 42°C, aliquots were serially diluted and plated on selective agar to enumerate donor (LB + cefotaxime [CTX] 1 µg/mL), recipient (LB + CHL 32 µg/mL), and transconjugant (LB + CTX + CHL) populations. Conjugation rates (γ, in mL/(CFU·h)) were estimated using the population-dynamic ‘approximate extended Simonsen’ model (1), which accounts for differential growth rates (*Ψ*) of the donor (*D*), recipient (*R*), and transconjugant (*T*) populations [50,51]. To ensure the robustness of our experimental approach, conjugation rate estimates were confirmed to be highly reproducible across three independent biological replicates, with strong and significant correlations observed between each replicate pair (Pearson’s *R* > 0.88, *P* < 0.001 for all; Figure S12).

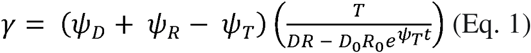

### Post-Conjugational Stability and Adjusted Rates

Plasmid stability was measured at 37°C using fresh transconjugants picked directly from selective mating plates. Picked colonies were grown overnight in non-selective LB. Given the high rates of instability observed in preliminary assays, loss was quantified between *t* = 0 and *t* = 1 hour by plating on selective (CTX) and non-selective LB. The plasmid loss rate (*ρ*, per hour) was calculated from the decline in the plasmid-carrying fraction (Equation 2).

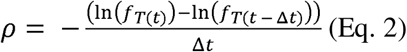

Loss-adjusted conjugation rates were then calculated by incorporating *ρ* into the Simonsen model (Equation 3). The fitness cost of plasmid carriage was quantified by comparing the maximum growth rates of transconjugant strains to their isogenic recipient parents in 96-well plates using an automated plate reader. Relative fitness was expressed as the ratio of the maximum growth rate of a transconjugant to its corresponding recipient.

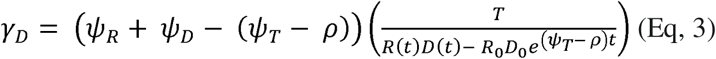

### Genomic Sequencing and Bioinformatic Analysis

Genomic DNA was extracted using the DNeasy Ultra Clean Microbial kit (Qiagen) for Illumina MiSeq sequencing (150 bp PE) and the Wizard Genomic DNA Purification kit (Promega) for Oxford Nanopore MinION sequencing (v14 chemistry, R10.4.1 flow cells; USEQ). Statistics are detailed in Tables S10 and S11. Reads were processed with Filtlong v0.2.1 (removing the bottom 5% and downsampling to 250 Mb) and Chopper v0.8.0 (-q 20, --minlength 500) to ensure high mapping quality. Donor and recipient hybrid assemblies were generated via Unicycler (v0.4.8) [52], while treanconjugants genomes were assembled using Flye (v2.9.2-b1786) [53] and polished with Medaka v1.8.0 [54]. All genomes were annotated using Bakta (v1.9.4) [55].

Genetic relatedness was assessed using PopPUNK (v2.4.0) [56]. Plasmidome and plasmid distances were calculated using Mash (v2.2.2; sketch 1000, k-mer 21) [57], with relatedness categorized as ‘highly related’ (Mash distance < 0.01), ‘moderately related’ (0.01 ≤ Mash ≤ 0.05), or ‘distantly related’ (> 0.05). Defense systems were identified using DefenseFinder (v1.3.0) [58]; individual gene hits were manually curated and collapsed into functional operons or anti-restriction clusters to prevent overestimating system abundance in multigenic systems.

For mutational analysis, 31 tranconjugant-recipient combinations (13 human: 7 *D*1, 6 *D*2; 18 chicken: 9 *D*1, 9 *D*2) were retained for high-resolution SNP analysis. These were selected from 39 initial tranconjugant isolates after excluding four sequencing failures and three strains with discordant sequencing pipelines that inflated mutation counts. Large-scale structural deletions were assessed across a broader set of 34 successfully assembled tranconjugant genomes (*D*1: 9 human, 9 chicken; *D*2: 8 human, 8 chicken).

To identify mutations, tranconjugant long-read genomes were compared to an artificial reference consisting of the parental recipient genome plus the donor plasmid. SNPs and small indels were called using breseq (v0.35.0) [59], while structural variants (>50 bp) were identified using SVIM (v1.4.2) [60]. Variants were filtered to exclude (i) pre-existing variants shared across transconjugants from the same recipient and (ii) technical artifacts, defined as high-frequency (>50%) mutations with >2 events per gene (indicative of read mismapping or multicopy genes). Major deletions were cross-referenced with insertion sequences identified by ISfinder [61] to investigate structural hotspots. This comprehensive pipeline captures the full spectrum of post-conjugational change, from single nucleotide variants to major adaptive rearrangements.

### Statistical Analysis

All statistical analyses were conducted in R v.4.4.0. A nested analysis of variance (ANOVA) was used to analyse the contributions of fixed effects (donor, temperature, origin) and random effects (strain nested within origin) on transfer rates, fitness, and loss rates. To meet the assumptions of normality and homoscedasticity, transfer rates were log10-transformed prior to analysis. Residual distributions were verified using Shapiro-Wilk tests and visual inspection of Q-Q plots. Relationships between continuous variables were assessed using Spearman’s rank correlation. *P*-values from multiple correlation tests were adjusted to control the False Discovery Rate (FDR) using the Benjamini-Hochberg method. Non-parametric comparisons were made using the Wilcoxon rank-sum test. To account for the discrete nature of count data, mutational patterns and plasmidomes defence/antidefense profiles were analysed using Poisson Generalized Linear Models (GLMs). This framework was applied to both total mutation frequencies and functional chromosomal categories. Statistical significance was defined as *P* < 0.05 unless otherwise specified.

## Author Contributions

M.B. and A.d.V. conceived and designed the study. M.B. performed all experiments, conducted the formal data analysis, and wrote the original draft of the manuscript. A.d.V. provided primary supervision for both the experimental work and the writing process. R.J.W. contributed to conceptualization and oversaw the writing and revision. E.F. developed the mathematical methodology for the conjugation and loss rate models and reviewed the manuscript. J.d.l.F.H. and A.S.M. provided supervision for the plasmid curing experiments and contributed to manuscript review and editing. A.S. secured funding and assisted with manuscript review. All authors read and approved the final version of the manuscript.

## Acknowledgments

This research was funded by the collaboration project ‘DiSSeMINATE’ (LSHM19183), co-funded by the PPP Allowance made available by Health∼Holland, Top Sector Life Sciences & Health, to stimulate public-private partnerships.

## Data Availability

Sequencing data have been made available on the European Nucleotide Archive under project PRJEB100537.

## Code Availability

All analysis scripts used in this study are available at the GitLab repository: https://gitlab.com/MatteoBuffoni/zoonotic_inci1-blactxm-1

## References

1. Murray, C. J. L. et al. Global burden of bacterial antimicrobial resistance in 2019: a systematic analysis. Lancet 399, 629–655 (2022).

2. Cassini, A. et al. Attributable deaths and disability-adjusted life-years caused by infections with antibiotic-resistant bacteria in the EU and the European Economic Area in 2015: a population-level modelling analysis. Lancet Infect. Dis. 19, 56–66 (2019).

3. Mendelson, M., Sharland, M. & Mpundu, M. Antibiotic resistance: calling time on the ‘silent pandemic’. JAC-Antimicrob. Resist. 4, dlac016 (2022).

4. De Angelis, G., Del Giacomo, P., Posteraro, B., Sanguinetti, M. & Tumbarello, M. Molecular mechanisms, epidemiology, and clinical importance of β-lactam resistance in Enterobacteriaceae. Int. J. Mol. Sci. 21, 5090 (2020).

5. Bush, K. & Jacoby, G. A. Updated functional classification of β-lactamases. Antimicrob. Agents Chemother. 54, 969–976 (2010).

6. Bevan, E. R., Jones, A. M. & Hawkey, P. M. Global epidemiology of CTX-M β-lactamases: temporal and geographical shifts in genotype. J. Antimicrob. Chemother. 72, 2145–2155 (2017).

7. Carattoli, A., Villa, L., Fortini, D. & García-Fernández, A. Contemporary IncI1 plasmids involved in the transmission and spread of antimicrobial resistance in Enterobacteriaceae. Plasmid 118, 102392 (2021).

8. De Been, M. et al. Dissemination of cephalosporin resistance genes between *Escherichia coli* strains from farm animals and humans by specific plasmid lineages. PLoS Genet. 10, e1004776 (2014).

9. Castañeda-Barba, S., Top, E.-M. & Stalder, T. Plasmids, a molecular cornerstone of antimicrobial resistance in the One Health era. Nat. Rev. Microbiol. 22, 18–32 (2024).

10. Hall, J. P. J., Brockhurst, M. A. & Harrison, E. Sampling the mobile gene pool: innovation via horizontal gene transfer in bacteria. Philos. Trans. R. Soc. B 372, 20160424 (2017).

11. Muloi, D. M. et al. Population genomics of *Escherichia coli* in livestock-keeping households across a rapidly developing urban landscape. Nat. Microbiol. 7, 581–589 (2022).

12. Kluytmans, J. A. J. W. et al. Extended-spectrum β-lactamase–producing *Escherichia coli* from retail chicken meat and humans: comparison of strains, plasmids, resistance genes, and virulence factors. Clin. Infect. Dis. 56, 478–487 (2013).

13. Liu, C. M. et al. *Escherichia coli* ST131-H22 as a foodborne uropathogen. mBio 9, e00470–18 (2018).

14. Van Hoek, A. H. A. M. et al. Transmission of ESBL-producing *Escherichia coli* between broilers and humans on broiler farms. J. Antimicrob. Chemother. 75, 543–549 (2020).

15. Madec, J.-Y., Haenni, M., Métayer, V., Saras, E. & Nicolas-Chanoine, M.-H. High prevalence of the animal-associated *bla*CTX-M-1 IncI1/ST3 plasmid in human *Escherichia coli* isolates. Antimicrob. Agents Chemother. 59, 5860–5861 (2015).

16. Valcek, A. et al. IncI1 ST3 and IncI1 ST7 plasmids from CTX-M-1-producing *Escherichia coli*obtained from patients with bloodstream infections are closely related to plasmids from *E. coli* of animal origin. J. Antimicrob. Chemother. 74, 2171–2175 (2019).

17. Kim, J. W. et al. Mechanisms of theta plasmid replication in enterobacteria and implications for adaptation to its host. EcoSal Plus 9, doi:10.1128/ecosalplus.ESP-0004-2019 (2020).

18. Kosterlitz, O. & Huisman, J. S. Guidelines for the estimation and reporting of plasmid conjugation rates. Plasmid 126, 102685 (2023).

19. Huisman, J. S. et al. Estimating plasmid conjugation rates: a new computational tool and a critical comparison of methods. Plasmid 121, 102627 (2022).

20. San Millan, A. Evolution of plasmid-mediated antibiotic resistance in the clinical context. Trends Microbiol. 26, 978–985 (2018).

21. Harrison, E., Guymer, D., Spiers, A. J., Paterson, S. & Brockhurst, M. A. Parallel compensatory evolution stabilizes plasmids across the parasitism-mutualism continuum. Curr. Biol. 25, 2034–2039 (2015).

22. Horesh, G. et al. A comprehensive and high-quality collection of *Escherichia coli* genomes and their genes. Microb. Genom. 7, 000499 (2021).

23. Alderliesten, J. B., Duxbury, S. J. N., Zwart, M. P., de Visser, J. A. G. M., Stegeman, A., & Fischer, E. A. J. (2020). Effect of donor-recipient relatedness on the plasmid conjugation frequency: a meta-analysis. BMC Microbiology, 20, 135.

24. Shaw, L. P., Rocha, E. P. C. & MacLean, R. C. Restriction-modification systems have shaped the evolution and distribution of plasmids across bacteria. Nucleic Acids Res. 51, 6806–6818 (2023).

25. Pinilla-Redondo, R. et al. Type IV CRISPR–Cas systems are highly diverse and involved in competition between plasmids. Nucleic Acids Res. 48, 2000–2012 (2020).

26. Pinilla-Redondo, R. et al. CRISPR-Cas systems are widespread accessory elements across bacterial and archaeal plasmids. Nucleic Acids Res.50, 4315–4332 (2022).

27. Benz, J. et al. Type IV-A3 CRISPR-Cas systems drive inter-plasmid warfare. ISME J. 17, 2221–2232 (2023).

28. Dimitriu, T. et al. Various plasmid strategies limit the effect of bacterial restriction–modification systems against conjugation. Nucleic Acids Res.52, 12976–12990 (2024).

29. Sengupta, M. & Austin, S. Prevalence and significance of plasmid maintenance functions in the virulence plasmids of pathogenic bacteria. Infect. Immun. 79, 2502–2509 (2011).

30. Baharoglu, Z., Bikard, D. & Mazel, D. Conjugative DNA transfer induces the bacterial SOS response and promotes antibiotic resistance development through integron activation. PLoS Genet. 6, e1001165 (2010).

31. Pons, M. C., Praud, K., Da Re, S., Cloeckaert, A. & Doublet, B. Conjugative IncC plasmid entry triggers the SOS response and promotes effective transfer of the integrative antibiotic resistance element SGI1. Microbiol. Spectr. 11, e02201–22 (2023).

32. Galhardo, R. S., Hastings, P. J. & Rosenberg, S. M. Mutation as a stress response and the regulation of evolvability. Crit. Rev. Biochem. Mol. Biol. 42, 399–435 (2007).

33. Souque, C., Escudero, J. A. & MacLean, R. C. Off-target integron activity leads to rapid plasmid compensatory evolution in response to antibiotic selection pressure. mBio 14, e02537–22 (2023).

34. Porse, A., Schønning, K., Munck, C. & Sommer, M. O. A. Survival and evolution of a large multidrug resistance plasmid in new clinical bacterial hosts. Mol. Biol. Evol. 33, 2860–2873 (2016).

35. Stearns, S. C. The evolutionary significance of phenotypic plasticity. BioScience 39, 436–445 (1989).

36. Björkman, J., Hughes, D. & Andersson, D. I. The cost of antibiotic resistance from a perspective of competing bacteria. Mol. Microbiol. 33, 265–273 (1999).

37. Patel, V., & Matange, N. (2021). Adaptation and compensation in a bacterial gene regulatory network evolving under antibiotic selection. eLife, 10, e70931.

38. Samuel, B., Mittelman, K., Croitoru, S. Y., Ben Haim, M. & Burstein, D. Diverse anti-defence systems are encoded in the leading region of plasmids. Nature 635, 186–192 (2024).

39. Kroll, J., Klinter, S., Schneider, C., Voss, I. & Steinbüchel, A. Plasmid addiction systems: perspectives and applications in biotechnology. Microb. Biotechnol. 3, 634–657 (2010).

40. Hall, J. P. J. et al. Plasmid fitness costs are caused by specific genetic conflicts enabling resolution by compensatory mutation. PLoS Biol. 19, e3001225 (2021).

41. San Millan, A., et al. Positive selection and compensatory adaptation interact to stabilize non-transmissible plasmids. Nat. Commun. 5, 5208 (2014).

42. Benz, F. & Hall, A. R. Host-specific plasmid evolution explains the variable spread of clinical antibiotic-resistance plasmids. Proc. Natl. Acad. Sci. U.S.A. 120, e2212147120 (2023).

43. Toribio-Celestino, L. et al. A plasmid-chromosome crosstalk in multidrug resistant enterobacteria. Nat. Commun. 15, 10859 (2024).

44. Alonso-del Valle, A., et al. Variability of plasmid fitness effects contributes to plasmid persistence in bacterial communities. Nat. Commun. 12, 2653 (2021).

45. Kosterlitz, O. & Huisman, J. Guidelines for the estimation and reporting of plasmid conjugation rates. Plasmid 100, 102685 (2023).

46. Dorado-García, A., et al. Molecular relatedness of ESBL/AmpC-producing *Escherichia coli* from humans, animals, food and the environment: a pooled analysis. J. Antimicrob. Chemother. 73, 339–347 (2018).

47. DelaFuente, J. et al. Within-patient evolution of plasmid-mediated antimicrobial resistance. *Nat*. Ecol. Evol. 6, 1980–1991 (2022).

48. Engler, C., Kandzia, R. & Marillonnet, S. A one pot, one step, precision cloning method with high throughput capability. PLoS ONE 3, e3647 (2008)

49. Pet zoldt, T. growthrates: Estimate growth rates from experimental data. R package (2022). URL: https://CRAN.R-project.org/package=growthrates

50. Simonsen, L., Gordon, D. M., Stewart, F. M. & Levin, B. R. Estimating the rate of plasmid transfer: an end-point method. J. Gen. Microbiol. 136, 2319–2325 (1990).

51. Huisman, J.-S., et al. Estimating plasmid conjugation rates: A new computational tool and a critical comparison of methods. Plasmid 121, 102627 (2022).

52. Wick, R. R., Judd, L. M., Gorrie, C. L. & Holt, K. E. Unicycler: resolving bacterial genome assemblies from short and long sequencing reads. PLoS Comput. Biol. 13, e1005595 (2017).

53. Kolmogorov, M., Yuan, J., Lin, Y. & Pevzner, P. A. Assembly of long, error-prone reads using repeat graphs. Nat. Biotechnol. 37, 540–546 (2019).

54. medaka: Sequence correction provided by ONT Research. https://github.com/nanoporetech/medaka, Accessed 22 Feb 2019.

55. Schwengers, O. et al. Bakta: rapid and standardized annotation of bacterial genomes via a curated sequence database. Microb. Genom. 7, 000685 (2021).

56. Lees, J. A., Harris, S. R., Parkhill, J. & Bentley, S. D. PopPUNK: a greedy, graph-based method for population definition from large-scale sequence data. Genome Res. 29, 304–316 (2019).

57. Ondov, B. D. et al. Mash: fast genome and metagenome distance estimation using MinHash. Genome Biol. 17, 132 (2016).

58. Tesson, F. et al. Systematic and quantitative view of the antiviral arsenal of prokaryotes. Nat. Commun. 13, 2561 (2022).

59. Deatherage, D. E. & Barrick, J. E. Identification of mutations in laboratory-evolved microbes from next-generation sequencing data using breseq. Methods Mol. Biol. 1151, 165–188 (2014).

60. Heller, D. & Vingron, M. SVIM: structural variant identification from long reads. Bioinformatics 35, 2907–2915 (2019).

61. Siguier, P., Perochon, J., Lestrade, L., Mahillon, J. & Chandler, M. ISfinder: the reference centre for bacterial insertion sequences. Nucleic Acids Res. 34, D32–D36 (2006).

