## Supplementary Material for "Post-transfer instability, not initial transfer, limits dissemination of IncI1-*bla*_CTX-M-1_ plasmids between chicken and human *Escherichia coli*"

**Supplementary Figures:**

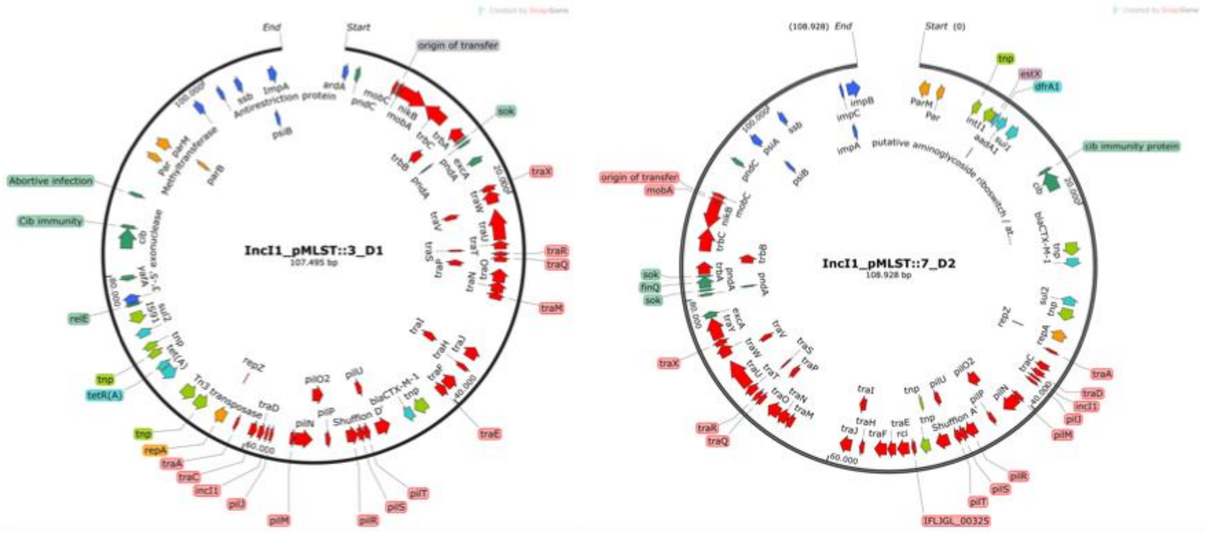


**Figure S1. Genetic maps of the IncI1-*bla*CTX-M-1 plasmids used in this study.** Circular maps represent the ST3 (left) and ST7 (right) lineages. Gene arrows are color-coded by biological function: red denotes conjugative transfer machinery; light blue indicates antibiotic resistance genes; dark green represents maintenance modules including toxin-antitoxin systems and partitioning machinery; blue indicates host-defense evasion and antirestriction genes; orange/yellow indicates replication (*rep*); and light green signifies genetic plasticity modules. Non-informative locus tags were removed, retaining only functionally descriptive gene names to emphasize biological roles and improve interpretability of the color-coded system.

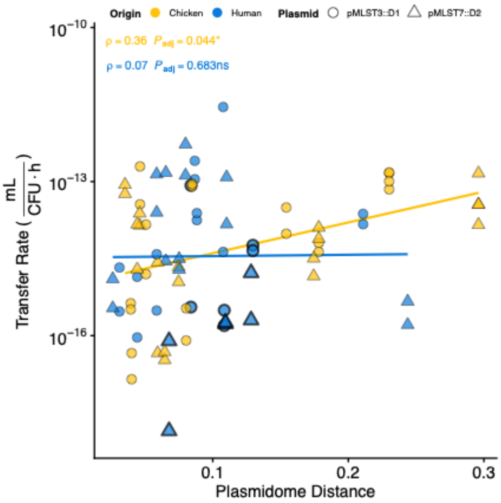


**Fig S2: Correlation between plasmidome distance and conjugation transfer rate.** Scatter plot showing the relationship between donor-recipient plasmidome distance and conjugation transfer rate (log10-scaled). Points are colored by recipient origin and shaped by donor plasmid. Plasmid-cured human strains are marked with a bold border. Spearman's rank correlation (ρ) and adjusted p-values are displayed for each origin group, with lines showing the linear regression fit.

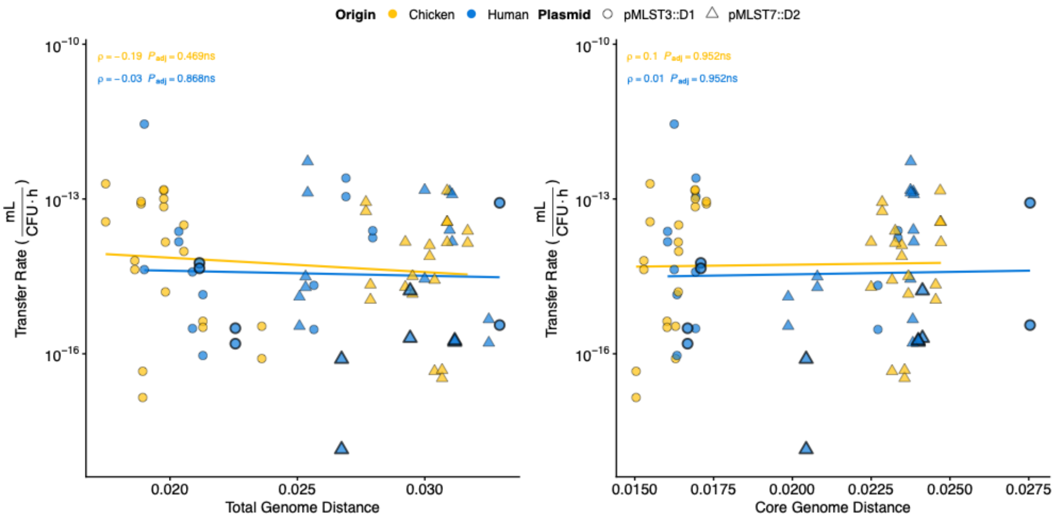


**Figure S3: Correlation of plasmid transfer rate with donor-recipient genomic distance.** Plasmid transfer rate is negatively correlated with core genome similarity and positively correlated with accessory genome content. Correlation between plasmid transfer rate and *D*-*R* recipient genome comparison. Panels show the relationship with (A) total genome size and (B) core genome size. Each point represents a single conjugation experiment, colored by the *R*'s origin (Human or Chicken) and shaped by the donor plasmid. Human-origin *R*s that were plasmid-cured prior to the experiment are highlighted with a thicker black border. Spearman's rank correlation (ρ) and Benjamini-Hochberg adjusted p-values are shown for each origin group.

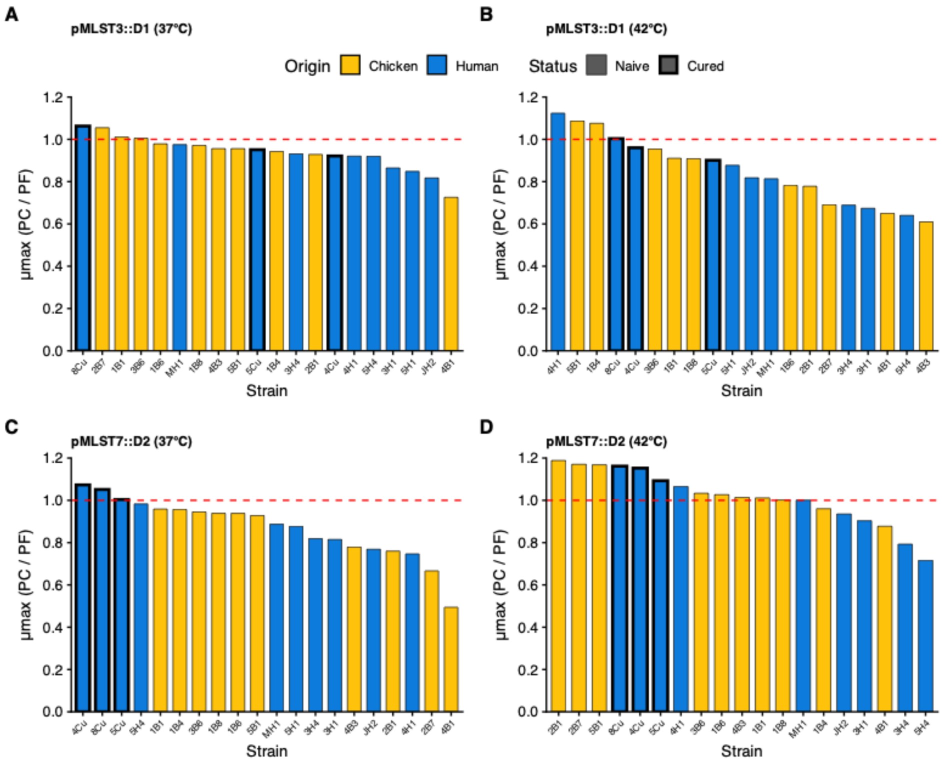


**Figure S4: Fitness effects of plasmid acquisition.** Bars represent the mean relative growth rate (µmax) of plasmid-carrying (PC) strains compared to their plasmid-free (PF) recipients, calculated from biological triplicates. Red dashed lines indicate a relative fitness of 1. Bold borders emphasize plasmid-cured strains.

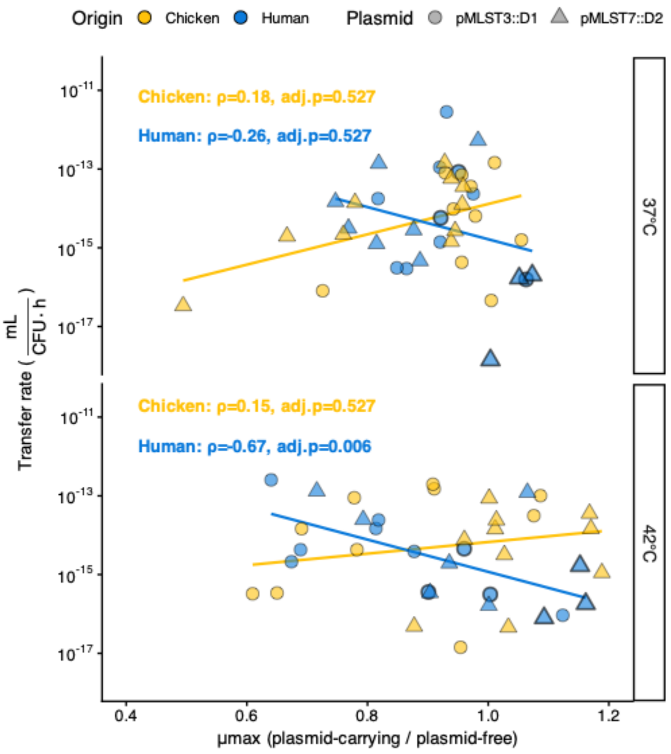


**Figure S5: Correlation between conjugation rate and relative growth rate.** Plasmid transfer rates (mL / CFU.h) are plotted against the mean relative growth rate (mu max) of transconjugants (PC/PF ratio, calculated from biological triplicates) at 37°C (top) and 42°C (bottom). Lines represent linear regression fits. Spearman's rank correlation coefficients (rho) are reported with Benjamini-Hochberg adjusted P-values (P-adj). Bold borders emphasize plasmid-cured strains.

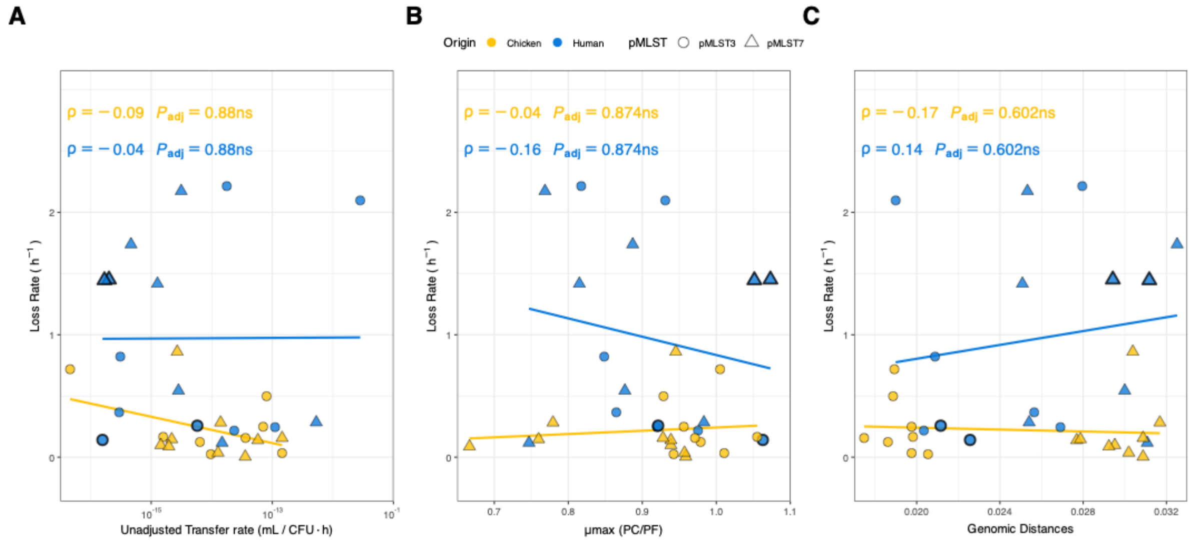


**Figure S6: Analysis of factors potentially influencing plasmid instability.** Panels A–C illustrate the relationship between Loss Rate ($h^{-1}$) and (A) Unadjusted Transfer Rates, (B) Relative Fitness, and (C) Genomic Distances, stratified by Origin (Chicken in yellow, Human in blue). Each point represents an observation; lines depict linear regression trends by group. Spearman's rank correlation coefficient (ρ) and adjusted p-values (Benjamini-Hochberg correction) are reported for each group. Significance level: *ns* = not significant ($p \geq 0.05$).

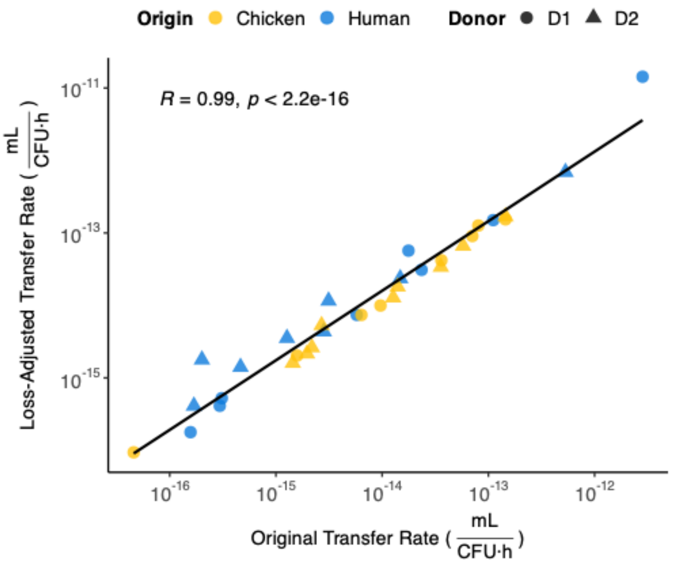


**Figure S7: Correlation between Original and Loss-Adjusted Transfer Rates.**
This figure illustrates the relationship between the original (raw) transfer rates and the transfer rates after adjusting for plasmid loss. A strong positive correlation (Spearman's ρ = 0.99) is observed, indicating that while adjusting for plasmid loss changes the absolute magnitude of transfer rates, it preserves the relative ranking and overall trends of transfer efficiency among the recipient strains.

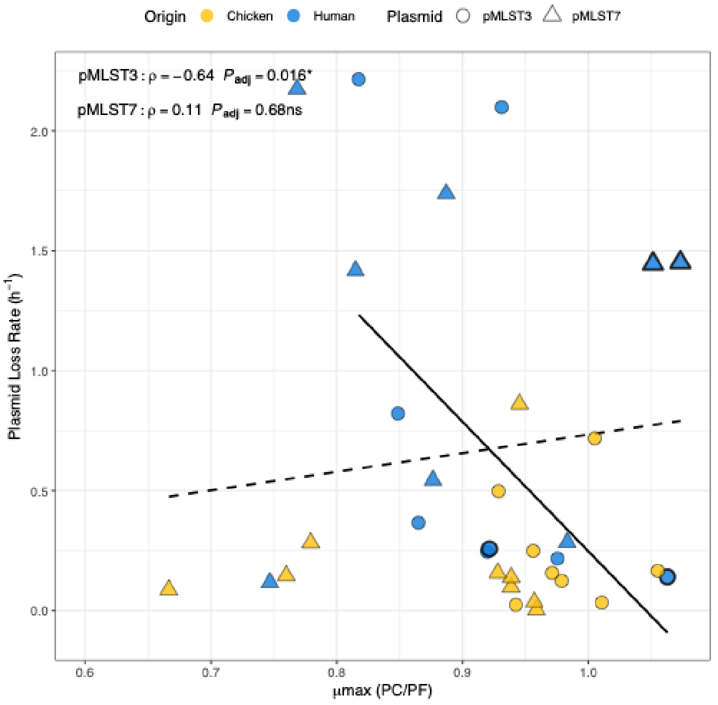


**Figure S8. Relationship between plasmid fitness cost and segregational stability is donor dependent.** The figure illustrates the relationship between the relative growth rate of transconjugants (T/R) and their plasmid loss rate ($h^{-1}$). Points are colored by the host origin of the recipient strain. The regression lines are stratified by donor plasmid. Spearman's rank correlation coefficient (ρ) and FDR-adjusted p-values, calculated separately for each donor group, are displayed in the top-left corner. Significance levels: * (p < 0.05), ns (not significant)

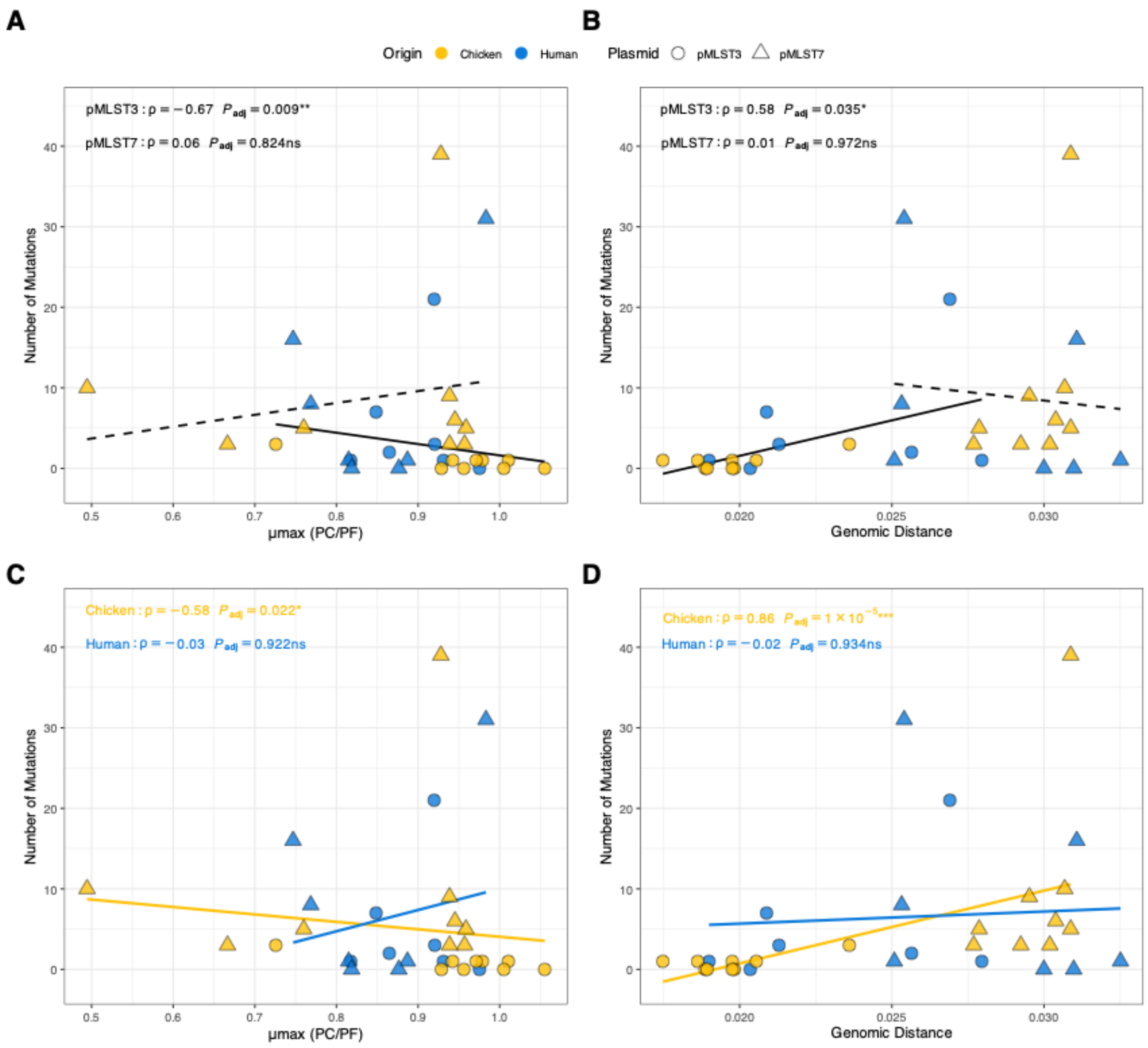


**Figure S9. Post-conjugational Mutational Dynamics by Donor Plasmid and Recipient Origin.** Each point represents a unique *T* (color: recipient origin, Chicken, yellow; Human, blue; shape: donor plasmid, pMLST::*D*1, circle; pMLST::*D*2, triangle). Panels A–B show correlations between Relative Fitness or Genomic Distance stratified by donor plasmid. Panels C–D show the same relationships stratified by recipient origin. Black or colored lines indicate linear regression trends. Spearman's $\rho$ and adjusted $p$-values (Benjamini–Hochberg FDR) are shown for each group.

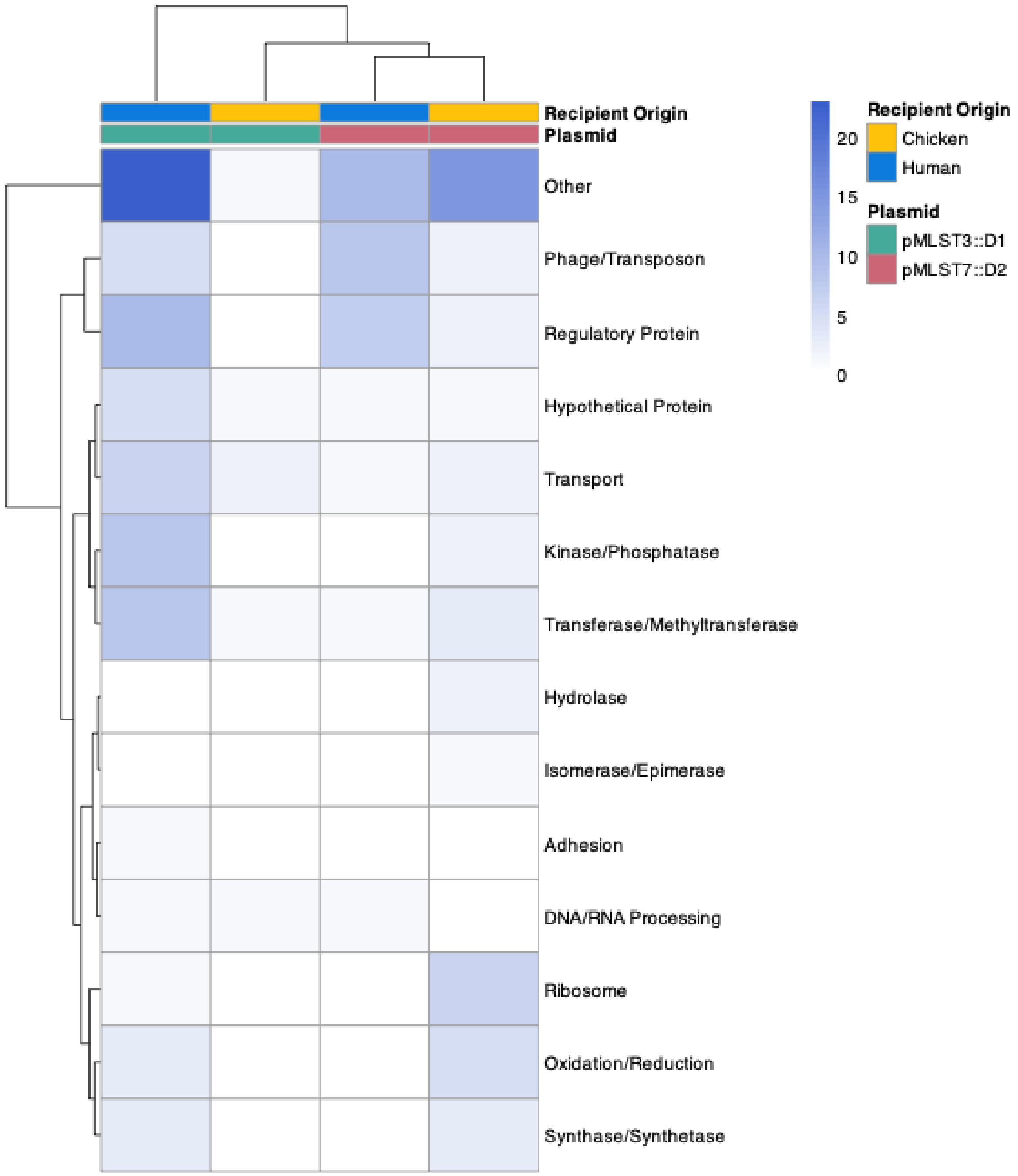


**Figure S10. Chromosomal mutation frequencies in transconjugants.**
Heatmap displays mutation frequencies in chromosomal gene categories (y-axis) across individual transconjugants. The top bar indicates recipient origin and donor plasmid. Blue shading intensity correlates with mutation counts.

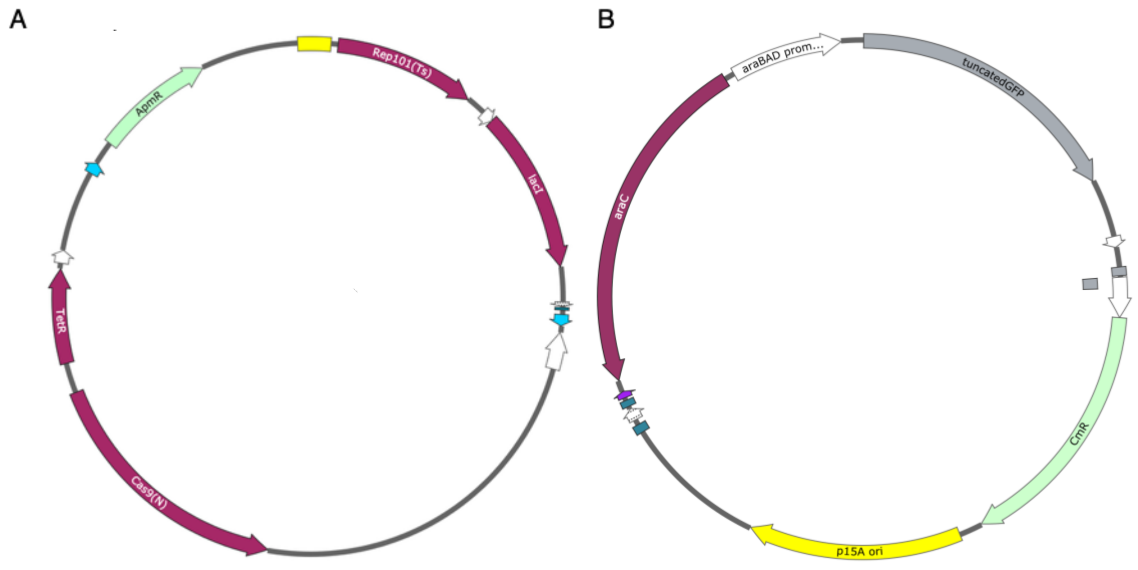


**Figure S11.** Graphical representation of the two plasmids used to cure the human *R* strains containing *bla*CTX-M-1 gene. A) pLC10 (10,548 bp) is the curing plasmid, which features a temperature-sensitive origin of replication (Rep101, yellow arrow) for controlled elimination. It carries the gene for the Cas9 nickase (Cas9, maroon arrow) to target the endogenous plasmid, as well as apramycin (ApmR, light green arrow) and tetracycline (TetR, maroon arrow) resistance genes for selection. B) pBC (4,102 bp) is a compatible reporter plasmid carrying a chloramphenicol resistance gene (CmR, light green arrow) for host strain selection and a p15A ori (yellow arrow). It also contains an arabinose-inducible system, consisting of the regulator araC (maroon arrow) and the araBADpromoter (white arrow), which controls the expression of a truncated GFP reporter gene (grey arrow).

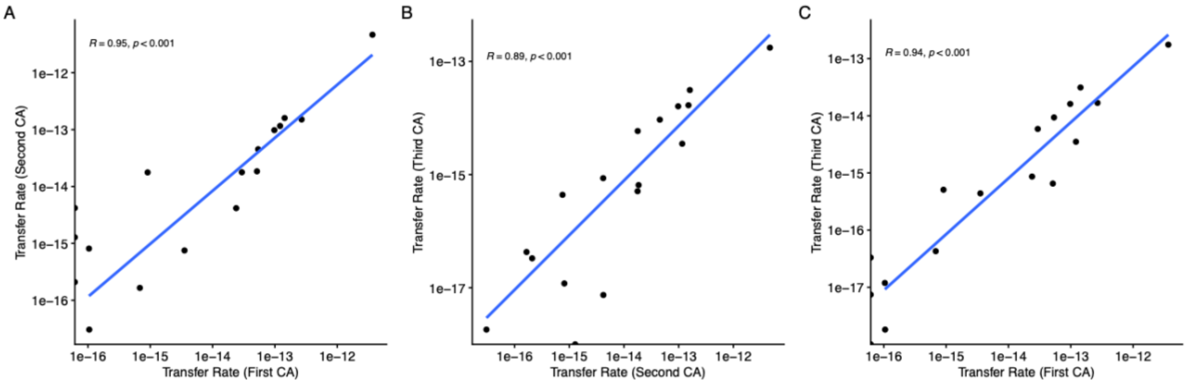


**Figure S12: High reproducibility of conjugation rate measurements across replicate experiments.** Scatter plots showing the correlation between transfer rate estimates (mL / (CFU·h)) from three independent biological replicates of the conjugation assay (CA). Each point represents the mean transfer rate for a single donor-recipient-temperature combination. (A) Correlation between the first and second replicate. (B) Correlation between the second and third replicate. (C) Correlation between the first and third replicate. Blue lines indicate the linear regression fit. Pearson's correlation coefficient (R) and the corresponding p-value are shown for each comparison. The axes are log10-scaled.

### Supplementary Tables

**Table S1.** Analysis of variance (ANOVA) of plasmid transfer rates.

| Source of Variation | Df[a] | SS[b] | MS[c] | F | P |
|---|---|---|---|---|---|
| Donor | 1 | 2.78E-25 | 2.78E-25 | 0.802 | 0.382 |
| Temp | 1 | 3.15E-25 | 3.15E-25 | 0.909 | 0.352 |
| Origin | 1 | 3.67E-25 | 3.67E-25 | 1.06 | 0.316 |
| Donor:Temp | 1 | 1.75E-25 | 1.75E-25 | 0.506 | 0.485 |
| Donor:Origin | 1 | 1.17E-25 | 1.17E-25 | 0.337 | 0.568 |
| Temp:Origin | 1 | 3.85E-25 | 3.85E-25 | 1.112 | 0.305 |
| Origin:Strain | 18 | 6.23E-24 | 3.46E-25 | 2.463 | 0.001** |
| Donor:Temp:Origin | 1 | 2.96E-25 | 2.96E-25 | 0.856 | 0.366 |
| Origin:Strain:Replicate | 20 | 1.63E-24 | 8.14E-26 | 0.235 | 0.633 |
| Residuals | 194 | 2.73E-23 | 1.41E-25 | | |

The table shows results from a mixed-effects ANOVA model where Strain was treated as a random factor nested within Origin.
[a] Df, Degrees of freedom; [b] SS, Sum of squares; [c] MS, Mean square.

**Table S2**. Analysis of variance (ANOVA) of plasmid fitness costs (i.e. relative fitness of *T*s).

| **Source of Variation** | **Df[a]** | **SS[b]** | **MS[c]** | **F** | **P** |
|---|---|---|---|---|---|
| Donor | 1 | 0.012 | 0.012 | 0.55 | 0.462 |
| Temp | 1 | 8.73E-05 | 8.73E-05 | 0.004 | 0.947 |
| Origin | 1 | 0.019 | 0.019 | 0.888 | 0.35 |
| Donor:Temp | 1 | 0.169 | 0.169 | 7.86 | 0.00701** |
| Donor:Origin | 1 | 0.006 | 0.006 | 0.294 | 0.59 |
| Temp:Origin | 1 | 0.046 | 0.046 | 2.136 | 0.15 |
| Origin:Strain | 18 | 0.929 | 0.052 | 2.4 | 0.00688** |
| Donor:Temp:Origin | 1 | 0.089 | 0.089 | 4.115 | 0.04744* |
| Residuals | 54 | 1.161 | 0.022 | | |

The table shows results from a mixed-effects ANOVA model where Strain was treated as a random factor nested within Origin.
[a] Df, Degrees of freedom; [b] SS, Sum of squares; [c] MS, Mean square

**Table S3.** Analysis of Variance (ANOVA) of Plasmid Loss Rates.

| **Source of Variation** | **Df[a]** | **SS[b]** | **MS[c]** | **F** | **P** |
|---|---|---|---|---|---|
| Origin | 1 | 4.615 | 4.615 | 34.455 | 5.50E-05*** |
| Donor | 1 | 0.179 | 0.179 | 1.338 | 0.2682 |
| Origin:Strain | 16 | 7.585 | 0.474 | 3.54 | 0.0132* |
| Residuals | 13 | 1.741 | 0.134 | | |

[a] Df, Degrees of freedom; [b] SS, Sum of squares; [c] MS, Mean square

### S4. Functional Annotation of Recipient's Plasmidome Defense and Anti-Defense Systems

| Strain | Origin | System Name | Category | Functional Annotation / Mechanism |
|---|---|---|---|---|
| **MH1** | Human | PD-T7-2 | Defense | Phage defense system specifically targeting T7-like phages. |
| **MH1** | Human | CBASS (Type III) | Defense | Cyclic-oligonucleotide-based signaling; triggers cell death to abort phage infection. |
| **MH1** | Human | DS-27 | Defense | Recently described phage defense system (mechanism under study). |
| **5H4** | Human | RM Type I | Defense | Classical restriction-modification; cleaves non-methylated foreign DNA. |
| **5H4** | Human | Gao_Ppl | Defense | Phage defense system utilizing specialized protein-protein interactions. |
| **5H1, 1B6** | Shared | AbiJ | Defense | Abortive Infection system: triggers programmed cell death upon phage entry. |
| **JH2, 3H1, 4H1, 5Cu** | Human | Mok_Hok_Sok | Defense | Toxin-Antitoxin (TA) system; provides population-level immunity and plasmid stability. |
| **4H1, 2B7, 3B6, 4B1** | Shared | MazEF | Defense | Type II TA system; induces reversible growth arrest or death under stress/infection. |
| **1B4** | Chicken | PrrC | Defense | Anticodon nuclease that cleaves host tRNA to block phage protein synthesis. |
| **1B4** | Chicken | ulx | Anti-Defense | Anti-Restriction protein; inhibits Type I Restriction-Modification enzymes. |
| **1B4** | Chicken | RM Type II | Defense | Simplest RM system; common barrier to horizontal gene transfer (HGT). |
| **2B7** | Chicken | RM Type III | Defense | Modification-dependent restriction; recognizes specific unmethylated motifs. |
| **2B7** | Chicken | Paris | Defense | Phage exclusion system that senses phage proteins and triggers cell arrest. |
| **2B7** | Chicken | AbiQ | Defense | Abortive infection system often found on mobile genetic elements. |
| **2B7** | Chicken | dar/ddr/hdf/ulx | Anti-Defense | Large multi-gene cluster providing high-level resistance to host restriction. |
| **3B6** | Chicken | TIR-III | Defense | Toll-interleukin receptor domain system; recognizes phage-induced stress. |
| **3B6** | Chicken | Pif | Defense | Phage exclusion system; prevents the replication of specific bacteriophages. |
| **3B6, 4B1, 4B3** | Chicken | ardb/psia/psib | Anti-Defense | Anti-Restriction (ardb) and Anti-SOS (psia/b) cluster; aids plasmid survival. |
| **3B6** | Chicken | Belisama | Defense | Phage defense system often co-localized with RM or TA systems. |
| **4B1** | Chicken | aca5 | Anti-Defense | Anti-CRISPR protein; specifically inhibits the host Cas immune complex. |

**Table S5.** Annotation of genes within the common deletion region (nucleotide positions 100,693 – 107,495) of Plasmid *D*1.

| **Gene Name** | **Start** | **End** | **Product Description** | **Function** | **Notes** |
|---|---|---|---|---|---|
| Antirestriction protein | 100,700 | 101,125 | Antirestriction protein | Stress adaptation, plasmid maintenance | Fully within deleted region |
| *yubJ* | 101,172 | 101,594 | Uncharacterized protein YubJ | *Uncharacterized* | Fully within deleted region |
| *yffA* | 101,591 | 101,782 | YffA (Uncharacterized protein) | *Uncharacterized* | Fully within deleted region |
| *ssb* | 102,552 | 103,079 | Plasmid-derived single-strand DNA-binding protein | Stress adaptation, plasmid maintenance | Fully within deleted region |
| hypothetical protein | 103,137 | 103,370 | hypothetical protein | *Uncharacterized* | Fully within deleted region |
| *yubM* | 103,339 | 105,387 | Uncharacterized protein YubM | *Uncharacterized* | Fully within deleted region |
| *psiB* | 105,442 | 105,876 | Conjugation system SOS inhibitor PsiB | SOS response, plasmid maintenance | Fully within deleted region |
| *psiA* | 105,873 | 106,592 | Plasmid SOS inhibition protein A | SOS response, plasmid maintenance | Fully within deleted region |
| *ygaA* | 106,589 | 107,185 | YgaA (Uncharacterized protein) | *Uncharacterized* | Fully within deleted region |

**Table S6**. Annotation of genes within the common deletion region (nucleotide positions 82,127 – 95,929) of Plasmid *D*2.

| Gene Name | Start | End | Product Description | Function | Notes |
|---|---|---|---|---|---|
| ***sok*** | 82,070 | 82,227 | sok antitoxin (CssrC) | Toxin-antitoxin system | Partially deleted (region 82,127-82,227 lost) |
| ***trbA*** | 82,364 | 83,572 | IncI1-type conjugal transfer protein TrbA | Conjugation initiation, mobilization | Fully within deleted region |

| | | | | | |
|---|---|---|---|---|---|
| ***trbB*** | 83,591 | 84,661 | IncI1-type conjugal transfer protein TrbB | Conjugation, mobilization | Fully within deleted region |
| ***trbC*** | 84,654 | 86,945 | F-type conjugative transfer protein TrbC | Conjugation, mobilization | Fully within deleted region |
| ***nikB*** | 86,982 | 89,681 | IncI1-type relaxase NikB | Conjugation initiation, mobilization | Fully within deleted region |
| ***mobA* (*nikA*)** | 89,692 | 90,024 | IncI1-type relaxosome accessory protein NikA | Conjugation initiation, mobilization | Fully within deleted region (Product = NikA) |
| ***oriT*** | 90,051 | 90,143 | origin of transfer | Mobilization, conjugation initiation | Fully within deleted region |
| ***mobC*** | 90,252 | 90,593 | Mobilizationdinucleotide biosynthesis protein MobC | Mobilization (likely, despite product name) | Fully within deleted region |
| ***DUF4942*** | 90,678 | 91,526 | domain-cont protein | *Uncharacterized domain* | Fully within deleted region |
| ***XRE*** | 92,106 | 92,357 | XRE family transcription regulator | *Regulatory* | Fully within deleted region |
| ***ydgA*** | 92,388 | 92,558 | YdgA | *Uncharacterized* | Fully within deleted region |
| **WP_000038339.1 (transposase)** | 92,620 | 93,561 | Transposase | Mobilization (related) | Fully within deleted region |
| ***ydfb*** | 93,626 | 93,892 | protein | *(Uncharacterized protein)* | Fully within deleted region |
| ***pndC*** | 93,984 | 94,418 | PndC | Toxin-antitoxin system (likely part of) | Fully within deleted region |
| ***ardA*** | 95,147 | 95,647 | Antirestrictic protein ArdA | Plasmid maintenance (antirestriction) | Fully within deleted region |
| ***deaR*** | 95,665 | 95,703 | DeaR family transcription regulator | *Regulatory* | Partially deleted (region 95,665-95,929 lost) |

**Table S7.** Unique, Non-Conserved Deletions in Donor Plasmids from Sequenced Transconjugants.

| Plasmid | Strain | Origin | Deleted Region[a] | Affected Gene Functions (Specific to Adjusted Non-Common Region) [b] |
|---|---|---|---|---|
| **D1** | 3H1 | Human | 76,748-100,692 | Regulatory Proteins (putA1); Toxin-Antitoxin Systems & Plasmid Maintenance (RelE, ImpB, ImpA, ImpC); DNA Modification, Repair & Metabolism (3'-5' exonuclease, XerD, Methyltransferase); Colicin Production/Immunity (Colicin-Ib, Colicin-Ib immunity protein); Phage Defense/Abortive Infection (Abortive infection protein); Plasmid Replication, Stability & Partitioning (ParM, ParB); Chaperones (GroEL); Hypothetical, Uncharacterized, or DUF-containing Proteins (YadA, YaeA, YaeB, YafA, ProQ/FinO, hypothetical, YagA, YbaA, YdeA, YdfA, DUF4238, CDS10, CDS11, YedA, Plasmid partitioning/stability, YubB, YfbA, Cytoplasmic, DUF1380, YchA) |
| **D1** | 3H1 | Human | 1-8,547 | Regulatory Proteins (agaR, XRE family); Restriction-Modification & Antirestriction (ardA); Toxin-Antitoxin Systems & Plasmid Maintenance (pndC); DNA Modification, Repair & Metabolism (Transposase); Conjugative Transfer & Mobilization (mobC, oriT, mobA, nikB); Metabolic & Biosynthesis Proteins; Hypothetical, Uncharacterized, or DUF-containing Proteins (YDFB, YdgA, DUF4942) |
| **D1** | 4H | Human | 100,682-100,692 | No gene functions specific to this adjusted region (all original genes were in the common D1 deleted region) |
| **D1** | 4H | Human | 5,036-5,164 | Hypothetical, Uncharacterized, or DUF-containing Proteins (DUF4942) |
| **D1** | JH2 | Human | 57,448-86,125 | Conjugative Transfer & Mobilization (incI1, TraD, TraC, TraA); Plasmid Replication, Stability & Partitioning (RepZ, RepA); DNA Modification, Repair & Metabolism (IS91 transposase, Tn3 transposase, IS6 transposase, IS110 transposase); Antibiotic Resistance (Tet(A), TetR(A), Sul2); Regulatory Proteins (putA1); Toxin-Antitoxin Systems & Plasmid Maintenance (RelE); Colicin Production/Immunity (YagA); Metabolic & Biosynthesis Proteins (Glm); Hypothetical, Uncharacterized, or DUF-containing Proteins (NusG-like, C4 RNA, Cytoplasmic, Drug/metabolite transporter, DUF4102, IS91 transposase, 3'-5' exonuclease, YadA, YaeA, YaeB, YafA, ProQ/FinO, hypothetical) |
| **D1** | 3B6 | Chicken | 1-18,007 | Regulatory Proteins (agaR, XRE family); Restriction-Modification & Antirestriction (ardA); Toxin-Antitoxin Systems & Plasmid Maintenance (pndC, sok, PndA, ArsR-gov); DNA Modification, Repair & Metabolism (Transposase); Conjugative Transfer & Mobilization (mobC, oriT, mobA, nikB, TrbC, TrbB, TrbA, ExcA, DotA/TraY); Metabolic & Biosynthesis Proteins; Hypothetical, Uncharacterized, or DUF-containing Proteins (YDFB, YdgA, DUF4942, HEAT repeat) |

| | | | | |
|---|---|---|---|---|
| **D1** | 3B6 | Chicken | 96,541-100,692 | Toxin-Antitoxin Systems & Plasmid Maintenance (ImpC, YubB); DNA Modification, Repair & Metabolism (YfbA, Methyltransferase); Restriction-Modification & Antirestriction (YchA); Hypothetical, Uncharacterized, or DUF-containing Proteins (Cytoplasmic, DUF1380) |
| **D2** | 1B1 | Chicken | 49,693-51,222 | Conjugative Transfer & Mobilization (Shufflon protein A'); DNA Modification, Repair & Metabolism (IS3 family IS1203 transposase ORF A, IS3 family IS629 transposase ORF B); Metabolic & Biosynthesis Proteins (AMP nucleosidase) |
| **D2** | 1B1 | Chicken | 51,373-51,464 | DNA Modification, Repair & Metabolism (IS3 family IS629 transposase ORF B) |
| **D2** | 1B1 | Chicken | 51,719-51,753 | Conjugative Transfer & Mobilization (Shufflon protein C) |
| **D2** | 1B8 | Chicken | 26,232-55,225 | DNA Modification, Repair & Metabolism (Transposase, IS91 transposase, IS3 IS1203, IS3 IS629, Shufflon-specific DNA recombinase); Regulatory Proteins (putA1); Metabolic & Biosynthesis Proteins (Glm, AMP nucleosidase); Antibiotic Resistance (Sul2); Plasmid Replication, Stability & Partitioning (RepA, RepZ); Conjugative Transfer & Mobilization (C4 RNA, TraA, TraC, TraD, IncI1, PilJ, Pili assembly chaperone, Type IVB prepilin, PilM, PilN, PilO2, PilP, Flp ATPase, PilR, PilS, PilT, PilU, Shufflon A', Shufflon C, TraE); Hypothetical, Uncharacterized, or DUF-containing Proteins (Transposase zinc-binding, NusG-like, (pseudo) hypothetical) |
| **D2** | 1B8 | Chicken | 96,009-102,602 | Toxin-Antitoxin Systems & Plasmid Maintenance (psiB, yubM, psiA); DNA Modification, Repair & Metabolism (ssb, yffA, yubJ); Restriction-Modification & Antirestriction (Antirestriction protein); Hypothetical, Uncharacterized, or DUF-containing Proteins (YgaA, DUF905) |
| **D2** | 2B1 | Chicken | 95,930-99,672 | Toxin-Antitoxin Systems & Plasmid Maintenance (FinQ, psiB, yubM, psiA); Hypothetical, Uncharacterized, or DUF-containing Proteins (YgaA) |
| **D2** | 4B3 | Chicken | 1-5600 | Plasmid Replication, Stability & Partitioning (ParM, Plasmid partitioning/stability, XerD, YedA); Chaperones (GroEL); DNA Modification, Repair & Metabolism (IS6 transposase); Hypothetical, Uncharacterized, or DUF-containing Proteins (CDS11, hypothetical) |
| **D2** | 4B3 | Chicken | 95,930-108,928 | Toxin-Antitoxin Systems & Plasmid Maintenance (FinQ, psiB, yubM, ImpC, ImpA, ImpB, psiA); DNA Modification, Repair & Metabolism (ssb, yffA, yubJ, Methyltransferase); Restriction-Modification & Antirestriction (Antirestriction protein, YchA); Hypothetical, Uncharacterized, or DUF-containing Proteins (YgaA, DUF905, DUF1380, Cytoplasmic, YubB ferredoxin-like) |
| **D2** | 3H1 | Human | 95,930-108,928 | Toxin-Antitoxin Systems & Plasmid Maintenance (FinQ, psiB, yubM, ImpC, ImpA, ImpB, psiA); DNA Modification, Repair & Metabolism (ssb, yffA, yubJ, Methyltransferase); Restriction- |

| | | | | |
|---|---|---|---|---|
| | | | | Modification & Antirestriction (Antirestriction protein, YchA); Hypothetical, Uncharacterized, or DUF-containing Proteins (YgaA, DUF905, DUF1380, Cytoplasmic, YubB ferredoxin-like) |
| **D2** | 3H4 | Human | 96,009-102,602 | Toxin-Antitoxin Systems & Plasmid Maintenance (psiB, yubM, psiA); DNA Modification, Repair & Metabolism (ssb, yffA, yubJ); Restriction-Modification & Antirestriction (Antirestriction protein); Hypothetical, Uncharacterized, or DUF-containing Proteins (YgaA, DUF905) |
| **D2** | 4H1 | Human | 1-6,698 | Plasmid Replication, Stability & Partitioning (ParM, Plasmid partitioning/stability, XerD, YedA); Chaperones (GroEL); DNA Modification, Repair & Metabolism (IS6 transposase); Hypothetical, Uncharacterized, or DUF-containing Proteins (CDS11, hypothetical) |
| **D2** | 4H1 | Human | 49,711-50,114 | Conjugative Transfer & Mobilization (Shufflon protein A'); DNA Modification, Repair & Metabolism (IS3 family IS1203 transposase ORF A) |
| **D2** | 4H1 | Human | 95,930-108,928 | Toxin-Antitoxin Systems & Plasmid Maintenance (FinQ, psiB, yubM, ImpC, ImpA, ImpB, psiA); DNA Modification, Repair & Metabolism (ssb, yffA, yubJ, Methyltransferase); Restriction-Modification & Antirestriction (Antirestriction protein, YchA); Hypothetical, Uncharacterized, or DUF-containing Proteins (YgaA, DUF905, DUF1380, Cytoplasmic, YubB ferredoxin-like) |
| **D2** | 5H4 | Human | 95,930-104,483 | Toxin-Antitoxin Systems & Plasmid Maintenance (FinQ, psiB, yubM, psiA); DNA Modification, Repair & Metabolism (ssb, yffA, yubJ); Restriction-Modification & Antirestriction (Antirestriction protein, YchA); Hypothetical, Uncharacterized, or DUF-containing Proteins (YgaA, DUF905, DUF1380) |
| **D2** | JH2 | Human | 5,877-6,698 | DNA Modification, Repair & Metabolism (IS6 family IS15 transposase) |
| **D2** | JH2 | Human | 52,379-82,126 | Conjugative Transfer & Mobilization (Shufflon-specific DNA recombinase, TraE, TraF, TraH, TraI, TraJ, TraM, TraN, TraO, TraP, TraQ, TraR, TraS, TraT, TraU, TraV, TraW, TraX, TraY, ExcA); DNA Modification, Repair & Metabolism (DNA primase, Putative endonuclease); Metabolic & Biosynthesis Proteins (Lipopolysaccharide core heptose(II)-phosphate phosphatase); Hypothetical, Uncharacterized, or DUF-containing Proteins ((pseudo) hypothetical, Conjugal transfer protein (multiple), ArsR-gov, PndA (multiple)) |
| **D2** | JH2 | Human | 95,930-108,928 | Toxin-Antitoxin Systems & Plasmid Maintenance (FinQ, psiB, yubM, ImpC, ImpA, ImpB, psiA); DNA Modification, Repair & Metabolism (ssb, yffA, yubJ, Methyltransferase); Restriction-Modification & Antirestriction (Antirestriction protein, YchA); Hypothetical, Uncharacterized, or DUF-containing Proteins (YgaA, DUF905, DUF1380, Cytoplasmic, YubB ferredoxin-like) |
| **D2** | MH1 | Human | 51,373-51,857 | DNA Modification, Repair & Metabolism (IS3 family IS629 transposase ORF B); Conjugative Transfer & Mobilization (Shufflon protein C) |

[a] Deletion coordinates are adjusted to specifically exclude predefined common conserved regions found in the parent plasmids (D1: 100,693-107,495; D2: 82,127-95,929).
[b] This column summarizes the biological roles of genes located only within the adjusted, non-common segments of the deletion. Functions of genes from the common conserved regions are excluded from this summary.

**Table S8**. Characteristics of the donor and cured strains and their *bla*$_{CTX-M-1}$ carrying-plasmids used in the study.

| Strain Names | Type | Plasmid Inc Type | pMLST | Plasmid Size (bp) | Comparison to *D*1 (Mash) [a] | Comparison to *D*2 (Mash) [a] | Sequence Shared with D1 (BLAST) [b] | Sequence Shared with D2 (BLAST) [b] |
|---|---|---|---|---|---|---|---|---|
| D1 | Donor | IncI1 | 3 | 107,495 | Full identity (0) [1000/1000] | Highly related (0.00763) [742/1000] | 100% | 92% |
| D2 | Donor | IncI1 | 7 | 108,928 | Highly related (0.00763) [742/1000] | Full identity (0) [1000/1000] | 91% | 100% |
| WT4 | Cured | IncI1 | 35 | 90,816 | Moderately related (0.0338) [326/1000] | Moderately related (0.0341) [323/1000] | 70% | 70% |
| WT5 | Cured | IncI1 | Unknown | 65,511 | Moderately related (0.0274) [391/1000] | Moderately related (0.0302) [361/1000] | 57% | 56% |
| WT8 | Cured | IncFII | 53 | 55,677 | Distantly related (0.1618) [17/1000] | Distantly related (0.1647) [16/1000] | 3% | 3% |

[a] Comparison to *D* plasmids *D*1 and *D*2 based on Mash distance. Qualitative descriptors ('highly', 'moderately', 'distantly' related) are based on thresholds defined in the Methods. The fraction in brackets represents the proportion of shared Mash hashes out of 1000.
[b] Percentage of the *D* plasmid sequence found in the query plasmid, determined by BLASTn alignment as described in Methods.

**Table S9.** Analysis of collateral plasmidome changes following curing of the *bla*$_{CTX-M-1}$-harboring plasmid.

| Strain | Plasmidome Size (bp) | Total Plasmids | Big Plasmids (>10kbp) | Small Plasmids (<10 kbp) | Collateral Change in Plasmidome (bp)[a] | Observed Collateral Events[b] |
|---|---|---|---|---|---|---|
| WT4 | 84,367 | 4 | 1 | 3 | — | — |
| WT4-Cured | 75,715 | 2 | 1 | 1 | 8,652 | Not Detected: Two plasmids (3,731 bp and 5,166 bp). |
| WT5 | 125,250 | 3 | 1 | 2 | — | — |

| | | | | | | |
|---|---|---|---|---|---|---|
| WT5-Cured | 116,923 | 1 | 1 | 0 | 8,327 | Integrated: One plasmid (2,703 bp). Not Detected: One plasmid (5,625 bp). |
| WT8 | 131,269 | 6 | 2 | 4 | — | — |
| WT8-Cured | 117,037 | 3 | 2 | 1 | 14,232 | Merged: Two plasmids (2,818 bp and 3,421 bp), resultant plasmid 5,319 bp. Not Detected: One plasmid (5,159 bp) and the separate circular form of an 8,169 bp plasmid. [c] |

[a] Net size difference of the non-target plasmidome between the wild-type and the cured strain assemblies.
[b] Summary of observed differences between the plasmidome assemblies of the parent (hybrid long/short-read assembly) and cured (long-read-only assembly) strains. Differences might be due to biological changes or technical issues.
[c] the net collateral change in WT8 results from the concurrent merger of two plasmids and the absence of detection of two others (one plasmid and one extrachromosomal form).

**Table S10:** Illumina Short-Read Sequencing and Mapping Statistics

| Strain ID | Total Input Reads | Total Input Bases (Mbp) | Mapped Paired Reads | % Mapped Paired Reads | Mapped Bases (Mbp) | Avg. Insert Size (bp) | Overall Base Error Rate (%) | Average Coverage (X) | % Ref. Bases Covered |
|---|---|---|---|---|---|---|---|---|---|
| **BR4(D1)** | 728742 | 125.82 | 360658 | 98.98 | 128.07 | 257.28 | 1.45 | 24.39 | 99.99 |
| **MH1** | 4457452 | 652.11 | 2200345 | 98.73 | 644.28 | 312.79 | 1.43 | 127.37 | 100 |
| **JH2** | 5425768 | 790.82 | 2678780 | 98.74 | 781.41 | 315.66 | 1.34 | 153.01 | 100 |
| **3H1** | 4298742 | 618.56 | 2119534 | 98.61 | 610.37 | 304.68 | 1.07 | 119.32 | 100 |
| **3H4** | 5491094 | 804.12 | 2708396 | 98.65 | 793.83 | 306.84 | 1.93 | 159.52 | 100 |
| **4H1** | 5143084 | 719.74 | 2530062 | 98.39 | 708.26 | 287.38 | 1.98 | 148.72 | 100 |
| **5H1** | 5618998 | 812.36 | 2764133 | 98.39 | 799.65 | 305.47 | 1.61 | 161.1 | 100 |
| **5H4** | 4816326 | 707.81 | 2366369 | 98.26 | 696.03 | 334.94 | 1.98 | 140.35 | 100 |
| **38.52(D2)** | 4248984 | 624.97 | 2093780 | 98.55 | 616.42 | 335.24 | 1.55 | 118.98 | 100 |
| **1B1** | 4529520 | 655.94 | 2234651 | 98.67 | 647.8 | 332.14 | 0.95 | 137.86 | 100 |

| | | | | | | | | | |
|---|---|---|---|---|---|---|---|---|---|
| **1B4** | 4359120 | 617.43 | 2133057 | 97.87 | 604.36 | 306.97 | 4.15 | 127.71 | 100 |
| **1B6** | 4349110 | 617.62 | 2152431 | 98.98 | 611.69 | 304.87 | 1.51 | 129.62 | 100 |
| **1B8** | 5160214 | 729.05 | 2556441 | 99.08 | 722.74 | 293.57 | 0.51 | 147.87 | 100 |
| **2B1** | 3153358 | 449.41 | 1558938 | 98.87 | 444.64 | 312.32 | 0.79 | 89.89 | 100 |
| **2B7** | 5593690 | 805.58 | 2732246 | 97.69 | 787.42 | 312.61 | 0.87 | 160.99 | 100 |
| **3B6** | 4886172 | 686.19 | 2406503 | 98.5 | 676.06 | 280.31 | 2.49 | 136.27 | 100 |
| **4B1** | 4692804 | 668.44 | 2282329 | 97.27 | 650.37 | 314.59 | 2.72 | 123.36 | 100 |
| **4B3** | 4882838 | 699.67 | 2365007 | 96.87 | 677.75 | 313.6 | 6.98 | 138.09 | 100 |
| **5B1** | 6174078 | 887.91 | 3011148 | 97.54 | 867 | 290.94 | 0.5 | 184.35 | 100 |

**Table S11:** Oxford Nanopore Long-Read Sequencing and Initial Assembly Metrics

| **Strain ID** | **Nr Scaffolds** | **Total Assembly Size (bp)** | **GC (%)** | **Max Scaffold Size (bp)** | **N50 (bp)** | **Nr Gaps** | **Total Reads (ONT)** | **% Mapped Reads (ONT)** | **Average Coverage (X, ONT)** |
|---|---|---|---|---|---|---|---|---|---|
| **WT4_Cu** | 3 | 5192708 | 50.81 | 5117104 | 5117104 | 0 | 46814 | 99.45 | 67.6 |
| **WT5_Cu** | 2 | 5217924 | 50.71 | 5101001 | 5101001 | 0 | 41180 | 99.59 | 67.63 |
| **WT8_Cu** | 4 | 5294317 | 50.78 | 5177280 | 5177280 | 0 | 46849 | 98.9 | 66.63 |
